# Plug-and-Play automated behavioral tracking of zebrafish larvae with DeepLabCut and SLEAP: pre-trained networks and datasets of annotated poses

**DOI:** 10.1101/2025.06.04.657938

**Authors:** Leandro A. Scholz, Tessa Mancienne, Sarah J. Stednitz, Ethan K. Scott, Conrad C.Y. Lee

## Abstract

Zebrafish are an important model system in behavioral neuroscience due to their rapid development and suite of distinct, innate behaviors. Quantifying many of these larval behaviors requires detailed tracking of eye and tail kinematics, which in turn demands imaging at high spatial and temporal resolution, ideally using semi or fully automated tracking methods for throughput efficiency. However, creating and validating accurate tracking models is time-consuming and labor intensive, with many research groups duplicating efforts on similar images. With the goal of developing a useful community resource, we trained pose estimation models using a diverse array of video parameters and a 15-keypoint pose model. We deliver an annotated dataset of free-swimming and head-embedded behavioral videos of larval zebrafish, along with four pose estimation networks from DeepLabCut and SLEAP (two variants of each). We also evaluated model performance across varying imaging conditions to guide users in optimizing their imaging setups. This resource will allow other researchers to skip the tedious and laborious training steps for setting up behavioral analyses, guide model selection for specific research needs, and provide ground truth data for benchmarking new tracking methods.

**SIGNIFICANCE STATEMENT:** Larval zebrafish are an emerging model in systems neuroscience, offering unique advantages for linking brain activity to behavior. However, detailed behavioral tracking, essential for such studies, requires time- and labor-intensive annotation and model training. To eliminate this bottleneck, here we provide a high-quality, annotated dataset of zebrafish behaviors for both free-swimming and head-embedded preparations alongside four pre-trained pose estimation models using DeepLabCut and SLEAP. We benchmark models’ performance across diverse imaging conditions to guide optimal setup choices. This community resource will allow researchers to bypass the most time-consuming stages of data annotation and training, enabling immediate behavioral analysis. By removing this key hurdle, this work will accelerate project initiation, support reproducibility, and provide a foundation for future tracking method development.

## INTRODUCTION

Tracking animal movements to quantify behavior accurately is essential to several fields of research, from biomechanics to neuroscience. For this reason, considerable efforts have been made to track animal behavior more precisely and with less labor-intensive methods (Berman, 2018; Pereira et al., 2020). To track animals in videos, it is necessary to process each frame to extract the position of the animal and other points or regions of interest using an image analysis workflow. This can be done in many ways, from a combination of classic image analysis steps such as background subtraction, image binarization and centroid finding (Burgess and Granato, 2007; Mirat et al., 2013; Marques et al., 2018; Guilbeault et al., 2021) to more complex methods such as model-based methods (Fontaine et al., 2008; Barreiros et al., 2021) and neural networks (Mathis et al., 2018; Pereira et al., 2019, 2022; Romero-Ferrero et al., 2019; Barreiros et al., 2021; Walter and Couzin, 2021; Gore et al., 2023).

In recent years, convolutional neural networks (CNNs) have enabled fully and semi-automated methods to track animal movement from images and can track animal poses with multiple body keypoints in the animal’s body, without requiring physical markers. Several groups have successfully implemented CNN architectures for the automated tracking of zebrafish in both 2 and 3 dimensions (Ravan et al., 2023; Fan et al., 2024), which may be challenging for beginners to implement compared to extensively documented methods such as DLC and SLEAP. While there is a large collection of pretrained markerless pose estimation networks for quadrupeds and mice (Ye et al., 2024), zebrafish data are not yet included in this repository. A few pre-trained models for zebrafish tracking with DLC exist, though they contain fewer keypoints or are trained on adult animals (Berg et al., 2022; Gore et al., 2023). Importantly, our annotations are highly detailed, and we provide a thorough comparison of DeepLabCut and SLEAP, which are among the most popular tools and are easily implemented into existing experimental frameworks (Lopes et al., 2015). However, these neural network based methods still require manually annotated images that reflect the range of movements and conditions encountered in an experiment. This is a time-consuming and labor-intensive process that requires hundreds of frames to build an accurate model. Quantifying the accuracy of any given model requires further post-hoc analysis, and some models may outperform others under different imaging conditions. Research groups often duplicate their efforts training models on similar images or select tracking methods that may underperform for their specific use case. Differences in analysis strategies also hinder data-sharing and direct comparisons between research groups, as outputs from different models can vary substantially depending on the number of keypoints and annotation strategy.

Here, we provide three large datasets of behavioral recordings and annotated pose images of zebrafish larvae (5-7 dpf), as well as pre-trained networks using open source methods DeepLabCut and SLEAP. We then quantify the performance of these networks under different imaging conditions and compare their relative strengths and weaknesses. The annotated dataset also serves as a ground truth to benchmark new models against existing methods. The pre-trained network can be used by other zebrafish researchers without need to annotate frames or as a starting point for their own networks, reducing the number of manual annotations that are necessary (Mathis et al., 2019). Lastly, we present results from an example analysis workflow, using the tracking output from the trained networks in an experiment where larvae were presented with visual and acoustic stimuli.

## METHODS

### Animals

All high-speed behavioral recordings were performed with approval from the University of Melbourne Office of Research Ethics and Integrity (in accordance with ethics approval 2022-24987-35220-5) and The University of Queensland Animal Welfare Unit (in accordance with ethics approval 2019/AE000341). Zebrafish (*Danio rerio*) larvae, of both sexes, were maintained at 28.5°C on a 14 h ON/10 h OFF light cycle. Adult fish were maintained, fed, and mated as previously described (Westerfield 2000). Experiments were carried out in larvae of the TL strain, mecp2^fh232^ or scn1lab^Δ44^ (Leong et al., 2015; Mendes et al., 2023; Wilde et al., 2024).

### Network Selection and Model Architectures

We chose to train networks using both DeepLabCut and SLEAP as they are maintained regularly, open source, well documented, and also popular within the field of neuroscience. For model architectures, we chose ResNet-50 and ResNet-152 (He et al., 2015) in DeepLabCut, where 50 and 152 relate to the number of layers in the network. This, in turn, is proportional to the number of parameters in the networks. Intuitively, one could expect that networks with the same base architecture, but more layers/parameters improve accuracy at the expense of processing time, with ResNet-50 processing videos faster than ResNet-152. For offline tracking, this is less of a concern. However, for applications requiring faster processing (at the cost of accuracy), such as real-time analysis, users may choose alternative networks available in DLC, such as MobileNet (Howard et al., 2017). With SLEAP, we chose to train the single animal (simple) and top-down methods, both built with the UNet architecture (Ronneberger et al., 2015). The simple model predicts keypoints for the entire image and groups into a single animal instance, whereas the top-down approach first locates a centroid to locate the animal then detects keypoints in the vicinity of the centroid found. Despite relying on two networks, the top-down method can outperform the single animal instance method in speed, as it first detects the animal then tracks keypoints in a smaller region of the image.

### Behavioral Datasets and Imaging Conditions

The workflow to obtain the trained models is depicted in Figure 1A. From a set of behavioral experiments, we obtained a series of videos of individual free-swimming larvae using our behavioral imaging apparatus previously described (Mancienne et al., 2021; Wilde et al., 2024; Favre-Bulle et al., 2025). The behavioral apparatus (Figure S1) featured a high-speed Ximea xiB-64 camera (model CB019MG-LX-X8G3, Ximea GmbH, Germany) equipped with a Sigma (Sigma Corporation, Japan) 17-70 mm f/2.8-4 lensor or a FLIR BlackFly BFS-U3-51S5M-C (Teledyne FLIR LLC, USA) camera and Tamron 10-40mm f1.4-inf lens (Tamron Co., Ltd, Japan), both equipped with infrared (IR) filters (530-750 nm).Visual stimuli were projected from an ultra-short throw ASUS (ASUSTeK Computer Inc., Taiwan)

**Figure 1:**
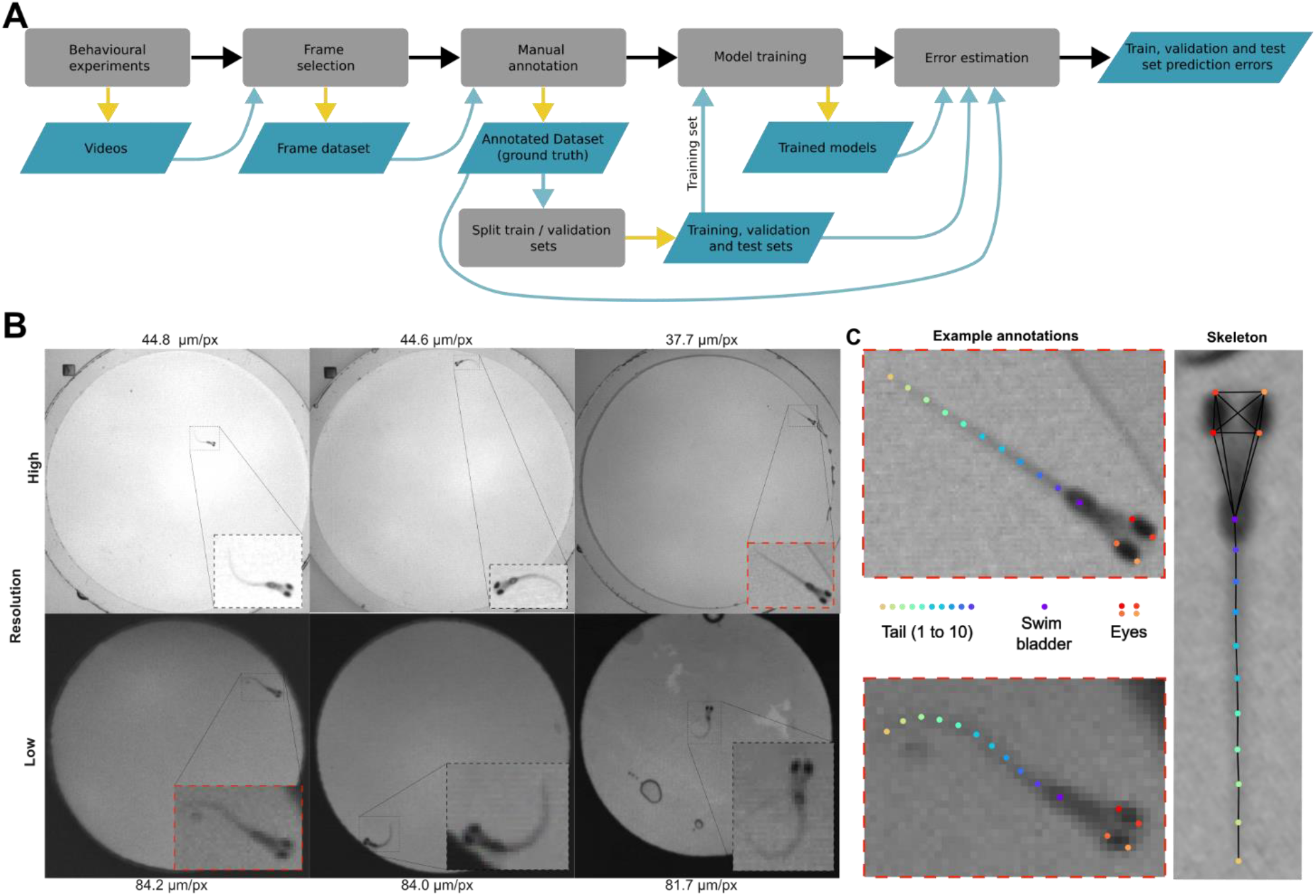
**A** Simplified block diagram describing the workflow to train and evaluate the pose estimation models. Blocks in gray represent steps in the process, whereas blocks in blue represent data (videos, images, spatial coordinates of annotations, etc..). Yellow arrows represent data outputs, and blue arrows input data. **B** Example of high (top row, 37-45 μm.px^-1^) and low resolution (81-88 μm.px^-1^) image frames from the dataset used to train and validate the pose estimation models with magnified regions of interest where fish were located (black and red dashed squares). **C** Expanded examples from two of the fish from **B** (red dashed lines) overlaid with our defined pose keypoints (10 keypoints in the tail, 1 in swim bladder and two for each eyes) and the connected skeleton (right).

P3E projector via a cold mirror (Edmund Optics, #64-442) onto a photography-grade Roscolux R116 Tough White (Rosco Laboratories Inc. USA) diffuser sheet attached to custom-made well plates using paraffin oil (Recochem Inc., Australia) to minimize optical distortions. IR illumination was provided by 850 nm LED strips (SMD5050-150, 30 LEDs/M) with a Roscolux R115 Light Tough Rolux (Rosco Laboratories Inc. USA) diffusing sheet. The behavioral arena (12.5 × 12.5 cm) was constructed from a transparent acrylic sheet, with interchangeable custom well plates made of acrylic and/or PDMS (*polydimethylsiloxane)*. Acoustic stimuli were delivered through speakers (Dayton Audio, USA) attached to the well plates and controlled by an amplifier (Dayton Audio, USA). All components, including the camera, projector, and amplifier, were controlled by a Lenovo P520 workstation (Intel® Xeon® W-2295 @ 3.00 GHz, 36 threads, 64 Gb RAM, Nvidia TITAN V), with the IR illumination and camera triggering managed by an Arduino UNO board. Experimental control and data acquisition were achieved using custom Bonsai-rx (Lopes et al., 2015) workflows, MATLAB (MathWorks, USA) scripts, or the Ximea CamTool software.

The experiments were performed under different imaging and spatial conditions (well diameters: 20 mm and 35 mm; camera exposure time: from 0.85 ms to 2 ms; gain: 6.02 dB and 12.04 dB) and various stimuli: spontaneous swimming, visual dark flash (300 Δlux), visual looms (R/V=500, initial loom size 0.9 mm, final loom size 25 mm) and acoustic stimulation (98 and 102 dB SPL). We then selected frames manually to ensure a variety of poses and image conditions (Figure 1B). To achieve a comprehensive dataset, the behavioral videos were acquired under different imaging conditions: low (81-88 μm.px^-1^) and high spatial resolution (37-45 μm.px^-1^), camera exposure times (0.7 to 3 ms), and contrasts (Figure 1B). Furthermore, we selected frames with spurious features that interfere with tracking. For example, frames where air bubbles were present or frames where parts of the body were occluded, as well as blurred tail points. These measures ensure the resulting network is applicable to a wide range of experimental conditions and is resilient to common sources of error.

To benchmark the models, we generated new datasets from videos acquired under specific camera exposure times and spatial resolutions. We refer to these as benchmarking datasets. The first benchmarking dataset was created from 5-6 dpf zebrafish larvae videos acquired using the same behavioral rig using 20 mm wells, where larvae experienced a 98 dB SPL tap, a 100 ms acoustic burst (1000 Hz) and a 1 s dark flash (300 Δlux). The conditions included four exposure times (0.5, 1, 3, and 10 ms) and four spatial resolutions (130-136, 97-103, 63-69, and 30-36 μm.px^-1^), thus resulting in 16 combinations of camera exposure and spatial resolution pairs. Each camera exposure time had a specific gain setup (0.5 ms with 18.06 dB, 1 ms with 12.04 dB and 3-10 ms with 0 dB). For each exposure/resolution pair, we had a set of unique videos, thus we called this dataset ‘independent’ (Figure 2). The second benchmarking dataset, which we call ‘downsampled’ (Figure 3), was generated to mitigate the effect of having unique videos for every exposure/resolution pair as well as the different gains. This dataset was created from the videos from the 30-36 μm/px resolution and downsampled to match the remaining three spatial resolutions.

**Figure 2:**
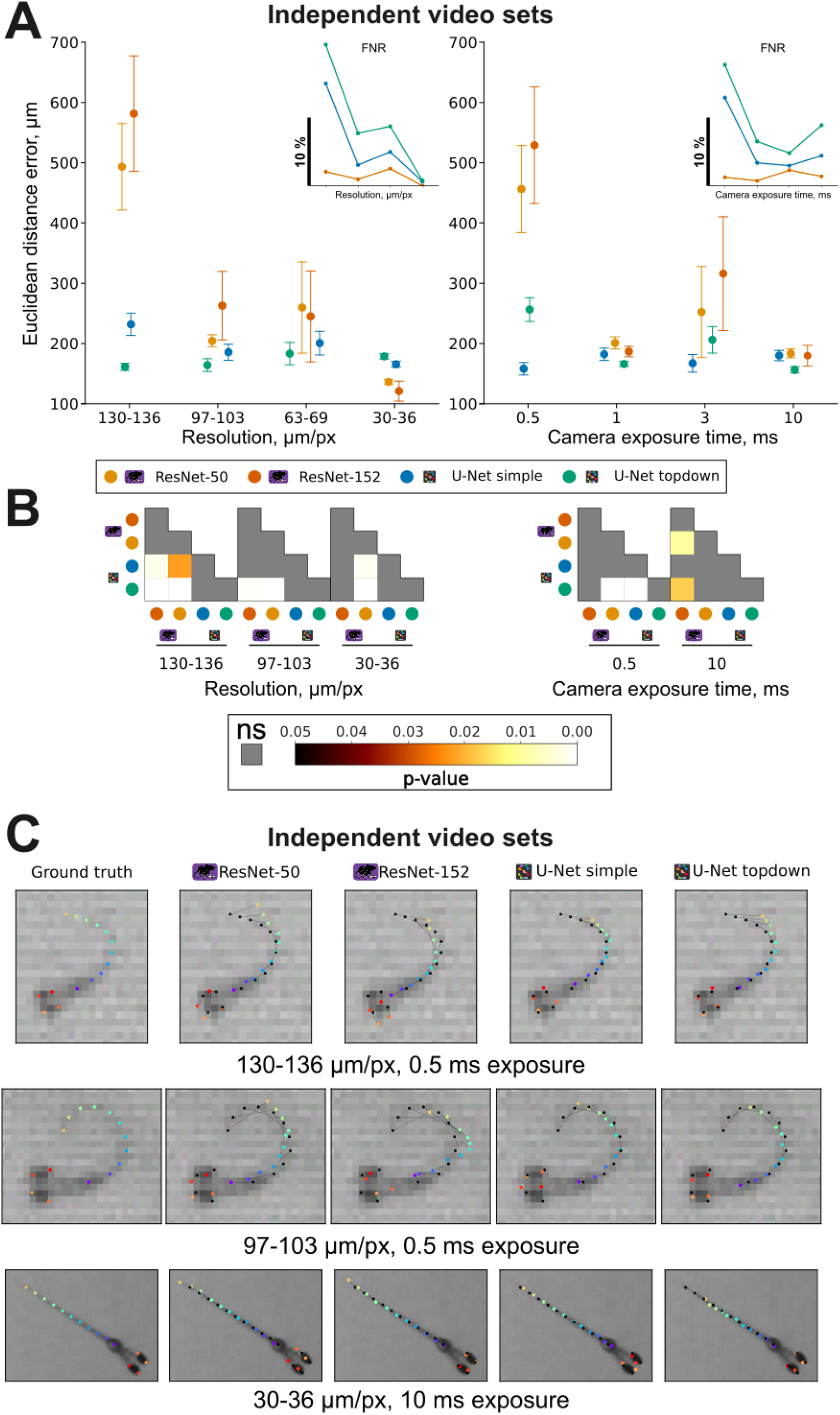
**A** Scatter plots of the average Euclidean distance error, in μm, for all pose keypoints grouped by resolution (left) and camera exposure (right) setting obtained by each pose estimation model (DeepLabCut Resnet-50 and Resnet-152 and SLEAP simple and topdown) using the independent benchmarking dataset. Inset scatter plots of false negative rate (FNR) grouped by spatial resolution (left) and camera exposure time (right) at the same order as the main plots. **B** p-value heatmaps for the most relevant pairwise comparisons of the Euclidean distance errors presented in **A**, obtained via Dunn test with Bonferroni correction for multiple comparisons (complete table available in Supplementary Table ST1). **C** Three example images (top from 130-136 μm.px^-1^, 0.5 ms camera exposure, middle, 97-103 μm.px^-1^, 0.5 ms exposure and bottom row 30-36 μm.px^-1^ and 10 ms exposure) with corresponding ground truth annotations (first column) and pose estimation network predictions (second to last columns). Colored dots in the model columns (second to last) are the actual model predictions and the black connected dots show the ground truth annotation.

**Figure 3:**
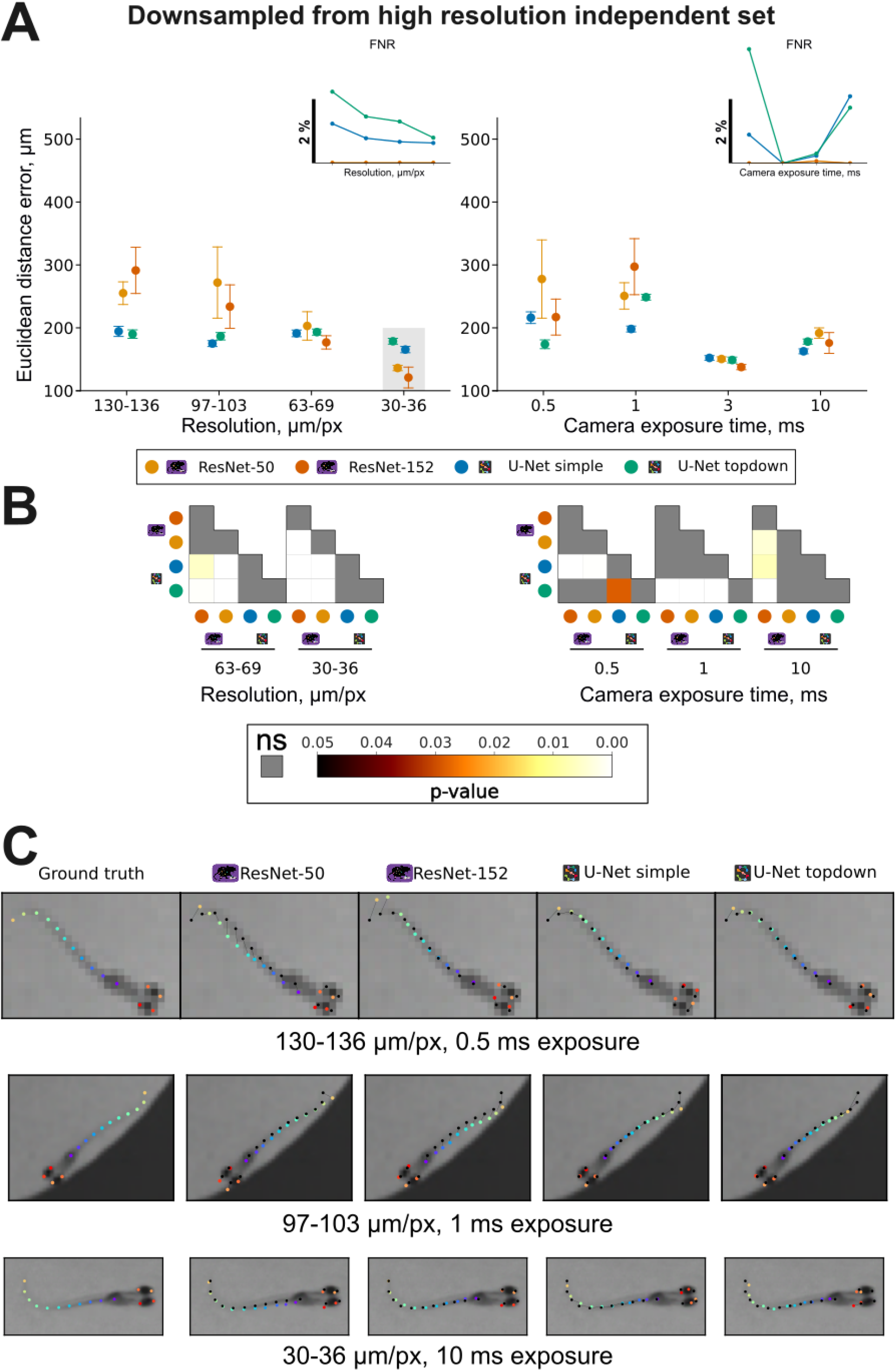
**A** Scatter plots of the average Euclidean distance error, in μm, for all pose keypoints grouped by resolution (left) and camera exposure (right) setting obtained by each pose estimation model (DeepLabCut Resnet-50 and Resnet-152 and SLEAP simple and topdown) using the downsampled benchmarking dataset. Gray shaded area in 30-36 μm.px^-1^ resolution results indicate that these data are the same as in Figure 2A, as these were the videos we used to generate the other lower resolution videos. Inset scatter plots of false negative rate (FNR) grouped by spatial resolution (left) and camera exposure time (right) at the same order as the main plots. **B** p-value heatmaps for the most relevant pairwise comparisons of the Euclidean distance errors presented in **A**, obtained via Dunn test with Bonferroni correction for multiple comparisons (complete table available in Supplementary Table ST1). **C** Three example images (top from 130-136 μm.px^-1^, 0.5 ms camera exposure, middle, 97-103 μm.px^-1^, 1 ms exposure and bottom row 30-36 μm.px^-1^ and 10 ms exposure) with corresponding ground truth annotations (first column) and pose estimation network predictions (second to last columns). Colored dots in the model columns (second to last) are the actual model predictions, and the black connected dots show the ground truth annotation.

Finally, we also provide an additional annotated dataset for the purpose of supporting benchmarking of new pose tracking methods. This dataset includes 12 videos of 5-7 dpf zebrafish larvae at various experimental conditions and a total of 512 annotated frames (see “additional annotated dataset” in our supplementary material).

### Network Training and Evaluation

We defined a zebrafish pose model that includes 15 keypoints (Figure 1C): 4 in the anterior and posterior edges of left and right eyes (2 points each), 1 point in the swim bladder (centroid), and 10 points throughout the tail. With the pose defined, we then annotated 28 behavioral recordings of free swimming and head-embedded animals for a total of 1641 frames. The complete list of parameters for each recording is available in the dataset description (supplementary material). Annotated frames were split into training and validation sets (90% training, 10% validation). To assess model generalizability, we also obtained another set of behavioral videos recorded under a wider range of resolutions, exposure times and camera gain setting. (Figure 2 and 3, and supplementary material).

To obtain the DeepLabCut models, we performed 5 training iterations, where we added new annotations to each iteration (up to 1641 images in the last set) to augment the dataset in cases where (i) we detected outlier frames using the DLC system and annotated a subset of them, or (ii) added frames from new videos that contained uncommon animal poses. To train the networks in SLEAP, we used the same dataset in a single iteration using all annotated frames. The evaluation results show that the DLC networks achieved train set errors from 2.46 to 1.00 pixels and validation set errors ranging from 3.41 to 1.29 pixels with ResNet-50 (full evaluation results in supplementary Figure S2A). With ResNet-152 train errors varied from 2.05 to 0.81 pixels, and validation sets from 2.94 to 1.05. Unlike DLC, we did not obtain evaluation results from intermediate network snapshots in SLEAP, but loss plots are presented in supplementary Figure S2B. SLEAP simple (single instance) achieved train and validation losses lower than 5.10^−5^ in 96 training iterations, whereas the top-down centered instance network achieved average losses lower than 0.002 for both training and validation sets in 69 iterations.

### Stimulus parameters and post-tracking analysis

Zebrafish larvae at 5-6 days post-fertilization (dpf) were placed in a custom well plate containing 7 individual wells. Each well was filled with 1 mL of E2 zebrafish medium. Larve were left in each well for 15 minutes prior to stimulus presentation to allow for acclimatization before presentation of either visual dark flashes (Δ 300 lux) or auditory taps (98 dB SPL). Each stimulus was repeated 5 times, and the order of presentation was randomized. Videos were recorded at 300 fps with a 0.5 ms exposure time, recording from –3 s to 3 s relative to stimulus onsets (total of 6 s) and had an inter-trial-interval of 1 minute.

Videos outputs contain images of 7 larvae in 7 wells and were segmented into 7 individual files for tracking, using opencv and ffmpeg. Once segmented into individual well videos, we could use DLC and SLEAP to track poses. The outputs from DLC and SLEAP produce x and y coordinates of each animal’s keypoint and the confidence of the estimation (labelled as “likelihood” in DLC, “instance” and “point” scores in SLEAP) for every frame in the video.

From the keypoints coordinates, two sets of metrics are calculated: point metrics and vector metrics. Point metrics are those that are calculated using a centroid calculated from a set of selected keypoints (e.g. eyes and swim bladder) or a single body keypoint (e.g. swim bladder). With this point, centroid position (in pixels, mm or normalized), distance travelled (in mm), velocity (mm.s^-1^), acceleration (mm.s^-2^) and distance from well center are calculated. Vector metrics are those that are calculated using at least two keypoints, which would define a vector. Among vector metrics, we also calculate mean tail curvature (degrees), tail velocity (degrees.s^-1^), tail acceleration (degrees.s^-2^), heading (degrees), heading turn angle (degrees). The vectors calculated are the heading vector and all tail segment vectors. The point and vector metrics are the basic metrics. With those, we can detect when swim bouts occurred. This is done using a peak finding function using smoothed distance travelled timeseries, along with a series of more complex conditions that include centroid acceleration, mean tail curvature data, and its rolling standard deviation timeseries as inputs. As the output of this ‘bout detector’, we obtained the frames where a bout started and ended (highlighted in Figure 4B and 4F).

**Figure 4:**
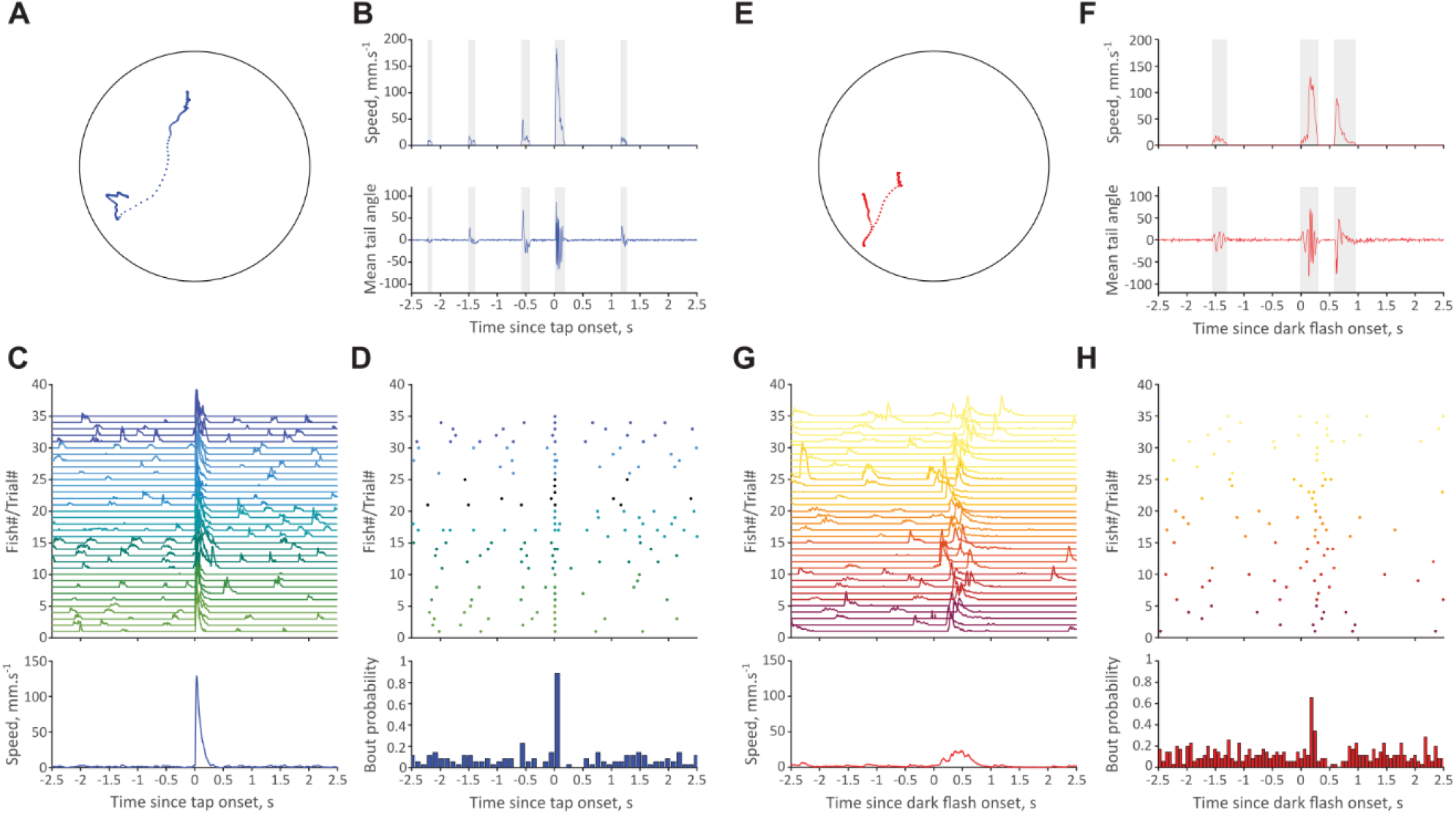
Example outputs from our analysis workflow using DLC prediction results as input. (**A, E**) Sample spatial trajectories within the 20 mm well for a single fish that experienced an acoustic tap (**A**) and a dark flash (**E**). Speed (top) and mean tail angle (bottom) timeseries data for the same example fish (**B, F**). Gray shaded areas show the intervals found by our bout detector. Speed time series results from multiple individual fish (7) trials (5) (top) and the average speed resulting from these trials (bottom) (**C, G**. Bout event plots for individual fish trials (**D, H**, top) and the binned bout probabilities resulting from these event plots (bottom).

## RESULTS

Accurate and efficient tracking of swimming behavior remains a key hurdle in zebrafish sensory neuroscience. To address this obstacle, we systematically evaluated the performance of two widely used pose estimation tools – DeepLabCut (DLC) and SLEAP, providing the community with a curated, annotated dataset of zebrafish larvae. We first used our benchmarking dataset and converted the errors, initially in pixels, to real distances, in μm to intuitively represent tracking accuracy across imaging conditions (Figure 2A, p-values within each group, 2B). The mean Euclidean distance errors including all key points were obtained using a small set of 1000 annotated frames from the videos of each exposure/resolution pair (approximately 60 per pair). Thus, slight variations in the error trends likely arise from the unique chosen frames having different predictability than frames from other exposure/resolution pairs. The average prediction error values of all trained networks (prediction errors of individual keypoints are presented in Supplementary Figures S4 to S17) ranged from 121 ± 17 μm (DLC-Resnet-152 at the highest image resolution, p<0.001 in any of the pairwise comparisons between DLC-Resnet-152 and any other models’ predictions at the same resolution, see Supplementary Table T1) to 581 ± 96 μm (DLC-Resnet-152 at the lowest image resolution).

Predictably, lower resolution images led to higher errors in both DLC and SLEAP models. DLC model predictions resulted in errors of 493 ± 72 μm for ResNet-50 and 581 ± 96 μm for ResNet-152. In contrast, both SLEAP model predictions were significantly more accurate than DLC (p=0.00199 SLEAP-simple vs. DLC-ResNet-152, p<0.0001 SLEAP-topdown vs. DLC-Resnet-152, p=0.002 SLEAP-simple vs. DLC-ResNet50, p<0.0001 SLEAP-topdown vs. DLC-Resnet-60). For high resolution videos, both SLEAP and DLC resulted in the lowest prediction errors, ranging from 100-200 μm. Although both models benefited from the increased resolution, DLC showed a greater improvement, with prediction errors decreasing at a greater rate with improved resolution. Specifically, the difference in average prediction errors between the highest and lowest resolutions was –357 μm for DLC-ResNet-50, –461μm for DLC-ResNet-152, –67 μm for SLEAP-simple and +18 μm for SLEAP-topdown. These results suggest that our DLC models are more sensitive to changes in image resolution, while SLEAP models maintain accurate predictions across different resolutions.

Model prediction errors as a function of camera exposure times varied from 156 ± 5 to 529 ± 97 μm (Mean ± SD), with DLC generally showing lower errors at higher exposure (p<0.0001 in pairwise comparisons between DLC-ResNet-152 at 0.5 ms vs. 1 ms, 3 ms and 10 ms, and p<0.0001 in comparisons between DLC-ResNet-50 at 0.5 ms vs. 3 ms and 10 ms, see Supplementary Table S1). In contrast, SLEAP prediction errors again were more consistent than DLC and no obvious trend was found. These results are more difficult to interpret, as the camera gain also varied for each exposure time in this dataset. However, the trend in the DLC results suggest that increasing exposure time may improve model performance in some cases. However, shorter exposures were paired with higher gain to maintain similar image dynamic range, which likely introduced additional image noise—potentially contributing to the higher prediction errors. In contrast, zebrafish larvae movements can often be extremely fast, with half tailbeats during a fast escape response lasting less than 10 ms. Fast tail movements can result in blurred images at camera exposure times greater than the movement durations, which will inevitably increase prediction errors. DLC networks varied more significantly with the changes in exposure time and gain. Low exposure times (0.5 ms), which also coincided with high gain setting, generated generally noisier images and resulted in higher prediction errors in DLC. The examples of predictions provided in Figure 2C and the figures S4 to S17 of the Euclidean distance prediction errors of individual keypoints suggest eye and the tail points further from the head are the main source of errors, but also show that overall, these predictions are sufficiently accurate to use for overall animal velocities and trajectories if tail kinematics are not essential.

Considering only tracking errors, SLEAP provides more consistent predictions that vary less with changes in resolution and exposure time. Despite this, for the false negative rate (FNR), which relates to the proportion of untracked keypoints (Figure 2A, inset plots), both SLEAP models have a greater FNR than DLC across all cases, with topdown showing higher FNR than the simple (single instance) model. More importantly, the FNR decreases significantly with increasing resolution, and varied from 20.66 to 0.79% in SLEAP topdown, from 15% to 0.63% in SLEAP simple. With varying exposure, the lowest FNR was found at 3 ms exposure (4.79% SLEAP topdown and 2.97 % SLEAP simple), and low exposure and noisy images (low exposure coupled with high gain) resulted in high FNR (17.74% SLEAP topdown and 12.91% SLEAP simple). At the same time, the FNR of both DLC models varied between 0.7% and 2.3%, with the exact same FNR in both networks. Thus, DLC and SLEAP operate by providing two distinct modes of failure: on one hand, DLC predicts body keypoints in most frames, but these may have low accuracy (false positives), whereas SLEAP may output a greater number of untracked points (false negatives), but an overall lower prediction error. The same way we mention that DLC prediction errors are more sensitive to changes in resolution, we can say that SLEAP FNR is also sensitive to changes in image resolution.

To minimize the confound of different gain values for each camera exposure time, we also generated a second dataset with constant gain values from the original test dataset by downsampling the highest resolution videos to obtain three sets of lower resolution videos (Figure 3A, p-values within each group in 3B). Moreover, the three example frame predictions in Figure 3C also show a clear difference in the noise levels from the independent dataset (Figure 2C). This second dataset also helps us address the fact that there are unique sets of frames for each resolution/exposure pair, as frames in each exposure time group are the same across all resolutions. Compared to the independent benchmarking dataset, the Euclidean distance prediction errors varied less, especially for the DLC networks, which ranged from 121 ± 1 (Resnet-152 at the 30-36 μm.px^-1^ resolution, p<0.0001 in each pairwise comparisons between the other models’ predictions at the same resolution) to 291 ± 37 μm.px^-1^ (Resnet-152 at the lowest resolution, 130-136 μm.px^-1^, mean ± SD). The Euclidean distance errors from SLEAP also varied less, from 149 ± 5 (top-down model at 3 ms exposure) to 249 ± 5 (top-down model at 1 ms exposure) μm. Within the two lower resolution sets (130-136 and 97-103 μm.px^-1^), no significant difference was observed between the models. With this dataset, we also find that the Euclidean distance errors may reach a minimum close to camera exposure times of 3 ms (p-values of pairwise comparisons in Table S1) and increase for the 10 ms exposure images. Finally, the example prediction results (Figure 3C, S4 to S17) corroborated the increased accuracy levels when noise is reduced in the videos as seen by the eyes and far tail points (tail 7 to 10) overall being closer to the ground-truth data.

To demonstrate the utility of the pre-trained models in an experimental setting, we tracked the behavior of seven 6 dpf zebrafish using DLC in responses to two commonly used sensory stimuli: an acoustic tap and a visual dark flash, with each fish receiving five presentations of each stimulus. For the acoustic tap, Figure 4A shows the spatial trajectory of an example fish, with Figure 4B showing its instantaneous swim speed and mean tail curvature aligned to stimulus onset. These traces capture both spontaneous bouts and a reliable, time-locked startle response marked by sharp increases in speed and tail bending. Figure 4C highlights the consistency of this response across animals and trials, showing robust startle response (increase in speed) following the tap. Figure 4D shows how speed and tail curvature can be used to identify bout onsets to quantify bout probability, which also rises sharply and consistently after the stimulus. Figures 4E–H show the same analyses for the visual dark flash. While fish still respond, the magnitude is lower, the latency is more variable, and both speed and bout probabilities reflect this increased temporal variability. Importantly, the tracking data produced from our DLC model effectively distinguishes subtle differences between types of startle responses arising from distinct neural circuits and allowing further in-depth bout classification based on tail kinematics (Burgess and Granato, 2007; Bhattacharyya et al., 2017; Marques et al., 2018; Jouary et al., 2024). Together, these data demonstrate how researchers can use our model to reliably track larvae and extract meaningful behavioral features across sensory modalities and experimental contexts, without the need for manual annotation.

## DISCUSSION

Zebrafish larvae at 5-7 dpf have a comparatively limited behavioral repertoire compared to adult animals, but the accurate quantification of these behaviors is nontrivial and an important task for researchers working in this model system across broad disciplines. Larval swim bout kinematics are complex and can only be obtained with high spatial and temporal resolution that most commercially available behavioral systems fail to capture. However, such high spatial and temporal resolution recordings generate large volumes of data that are impractical to annotate manually within a reasonable time, necessitating semi or fully automated methods. For example, annotating a single 300 fps 5 min duration video with 15 animal keypoints would take 250 hours, assuming an average of 10 seconds to annotate one frame. To overcomes this labor-intensity, our annotated dataset and pretrained models enable researchers to start tracking animals quickly, reliably, and easily. We also outline the specific impact of changing imaging parameters (e.g., camera gain, resolutions, exposure time, and lighting), allowing other users to replicate similar imaging conditions. In such cases, only a minimal number of frames need to be annotated to fine-tune our pre-trained model as a basis for their own custom model (Mathis et al., 2019). By making our dataset publicly available, we also offer a benchmarking tool for image analysis and computer vision developers aiming to create faster and more accurate methods to track zebrafish larvae behavior.

Unsurprisingly, we found that spatial resolution was the most important factor in tracking accuracy. Interestingly, in very low-resolution regimes, SLEAP outperformed DLC, indicating it may be a superior choice when other experimental constraints necessitate low spatial resolution (for example, high-throughput imaging with many animals). However, this improvement in accuracy comes with a trade-off: SLEAP tends to generate more false negatives, where keypoints are missed even if a centroid is detected. Therefore, the tolerance for missing data in an analysis should be carefully considered when selecting between these models.

We found that, while exposure time did not directly influence prediction errors, lower exposure time results in slightly lower contrast and subsequently higher prediction errors. Furthermore, tail kinematics of larval movements requires short camera exposure times (<3 ms), since some zebrafish larvae bout types can be extremely fast (e.g. short latency C-starts). Increased exposure times, despite increasing image contrast, also causes motion blur during these fast movements. Thus, to operate at low exposure times, it is important to adjust the strength of the IR illumination in the system to minimize its negative effect on image contrast. Ultimately, researchers need to balance their image quality needs within the constraints of their experiment, while optimizing image quality to capture the behaviors of interest.

The analysis workflow we present exemplifies how outputs of the pose estimation models can be analyzed but are certainly not limited to that. Once high-quality detailed zebrafish pose data are acquired with the pre-trained DLC and SLEAP networks, researchers can use one of the many existing analysis tools to further understand the neural and behavioral phenomena they study (Luxem et al., 2022; Schneider et al., 2023). One avenue we do not present in the current work but is increasingly used to understand zebrafish larval behaviors is the categorization of bout types, and further exploration of the spontaneous or stimulus-evoked behavioral sequence patterns (Marques et al., 2018; Johnson et al., 2020; Mearns et al., 2020; Reddy et al., 2020; Jouary et al., 2024).

Altogether, we provide a detailed, annotated dataset and pre-trained pose estimation networks for research groups studying larval zebrafish behavior. We further compared the performance of two popular network architectures (DeepLabCut and SLEAP) under variable imaging conditions. Although the network architectures we compare here are designed for offline analysis, it is feasible to use our annotations to produce speed-optimized models that permit real-time tracking. These resources will aid researchers by saving considerable time and helping them make informed selections to achieve the best performance for their experiments.

## DATA AVAILABILITY

Datasets, exported network weights DLC and SLEAP project files and code are available at https://github.com/Scott-Lab-QBI/zf_tracking_networks.

## AKNOWLEDGEMENTS

The authors would like to acknowledge The University of Queensland’s Biological Resources aquatics team and the Danio rerio University of Melbourne facility (DrUM, Melbourne, Australia) for maintenance of zebrafish lines.

## ETHICS

All work was performed in accordance with ethics approval from The University of Melbourne’s (2022-24987-35220-5) and The University of Queensland’s (2019/AE000341) Animal Welfare Units.

