## Supplementary Figures and Table for "Plug-and-Play automated behavioral tracking of zebrafish larvae with DeepLabCut and SLEAP: pre-trained networks and datasets of annotated poses"

1 – Department of Anatomy and Physiology, The University of Melbourne, VIC, Australia
2 – Queensland Brain Institute, The University of Queensland, QLD, Australia
* Corresponding author

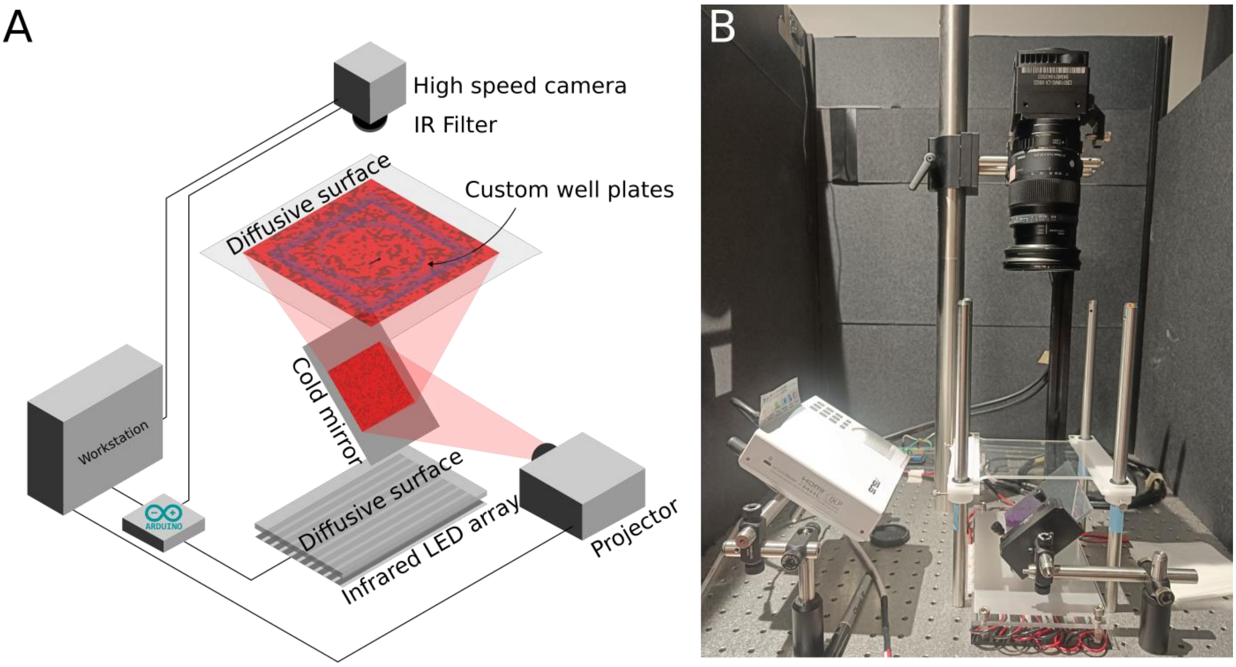


**Figure S1** – **A** Schematic representation of the behavioural apparatus. **B** Image of the behavioural apparatus installed.

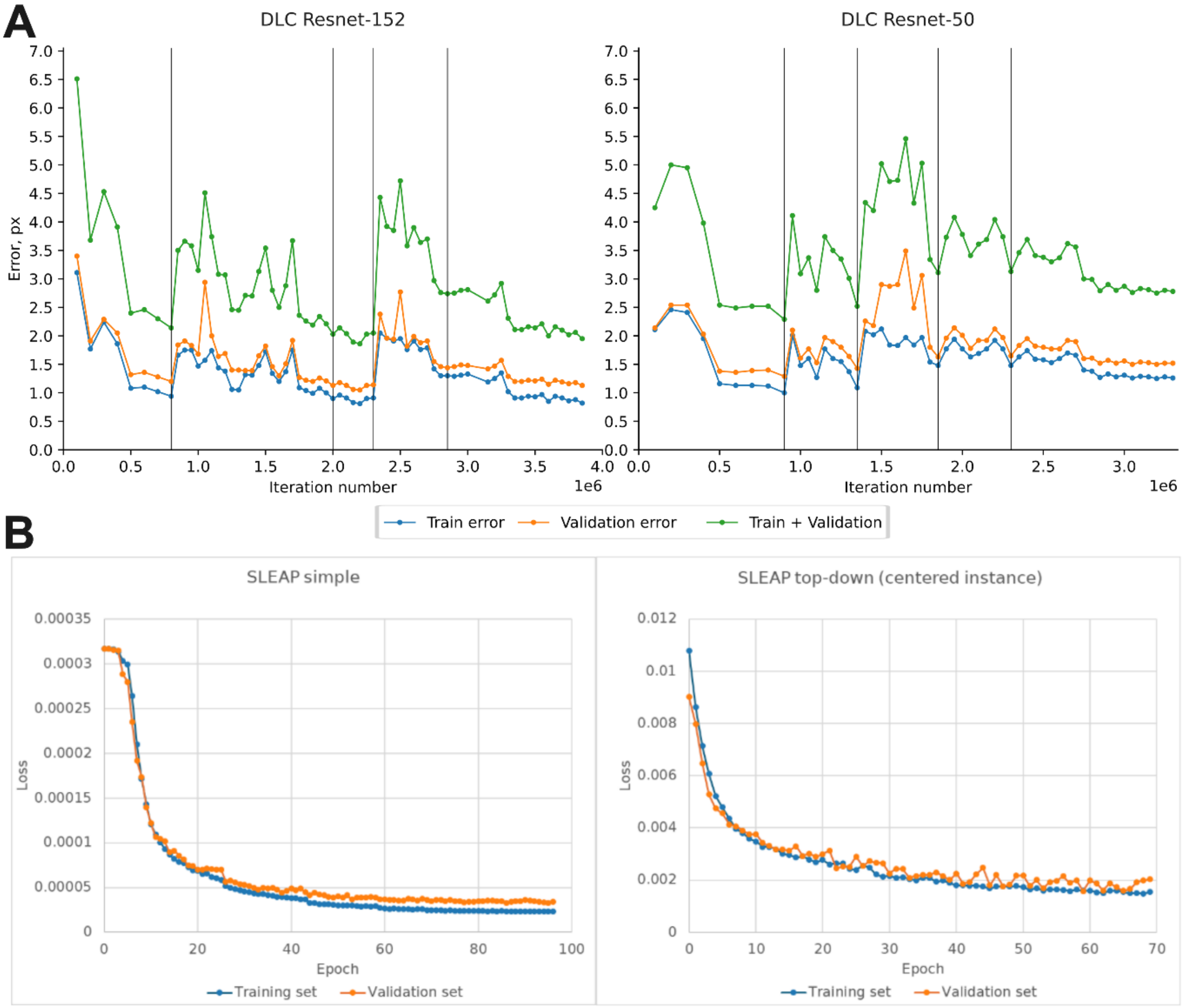


**Figure S2** – **A** Resnet-152 (left) and Resnet-50 (right) errors for the train (blue), validation (orange) and the sum of the train and validation (green) sets for each saved network snapshot during DeepLabCut training. Vertical lines indicate the iterations where new annotated frames were added to the train and validation sets. **B** SLEAP model train (blue) and validation (orange) set losses per epoch during training.

Figures S2 and S3 occupy multiple pages for ease of visualization and start in this page, with the captions shown at the end of each figure.

### Frame grid legend

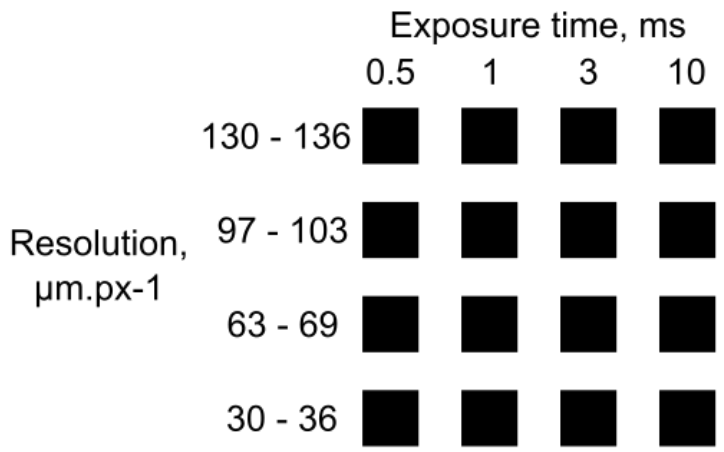


### Examples from the independent video set

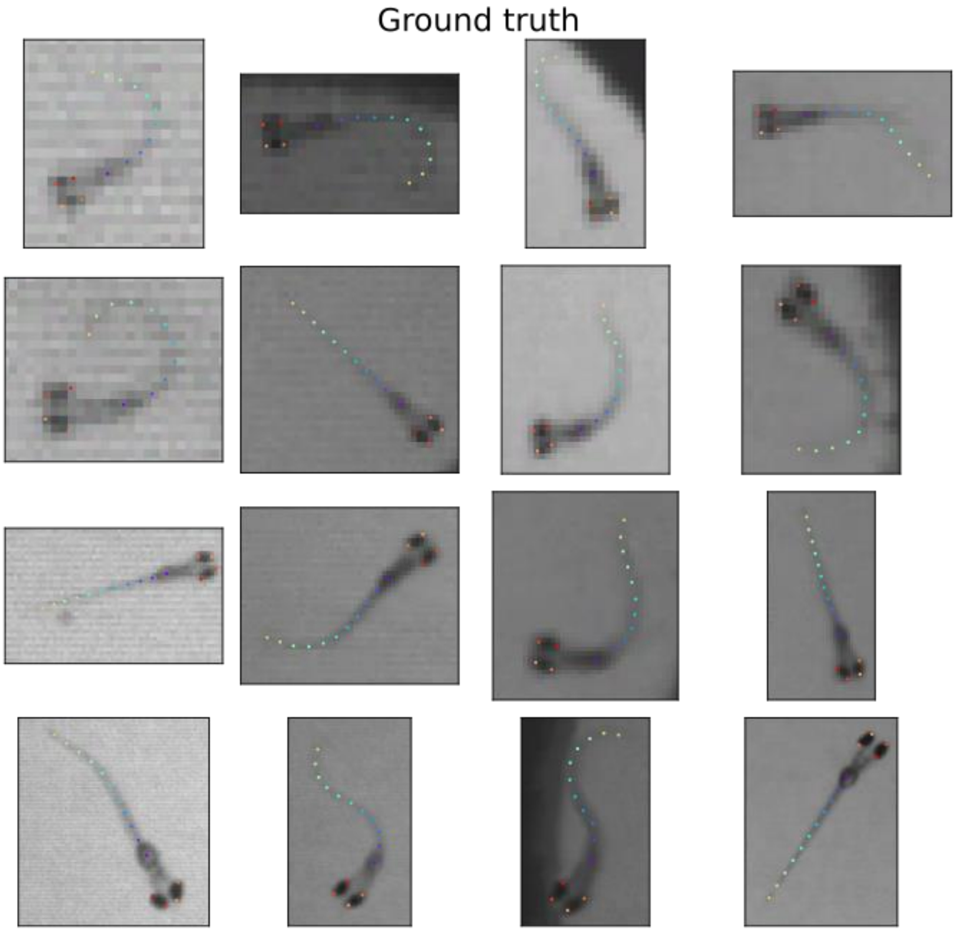

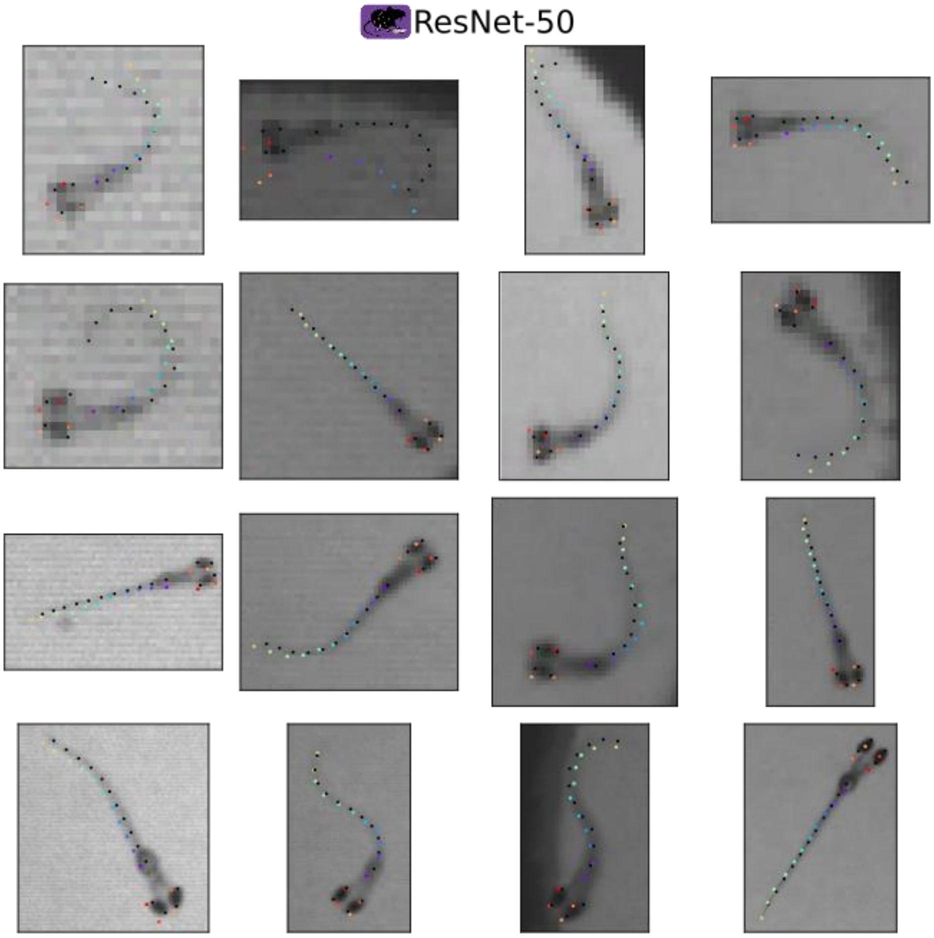
ResNet-50

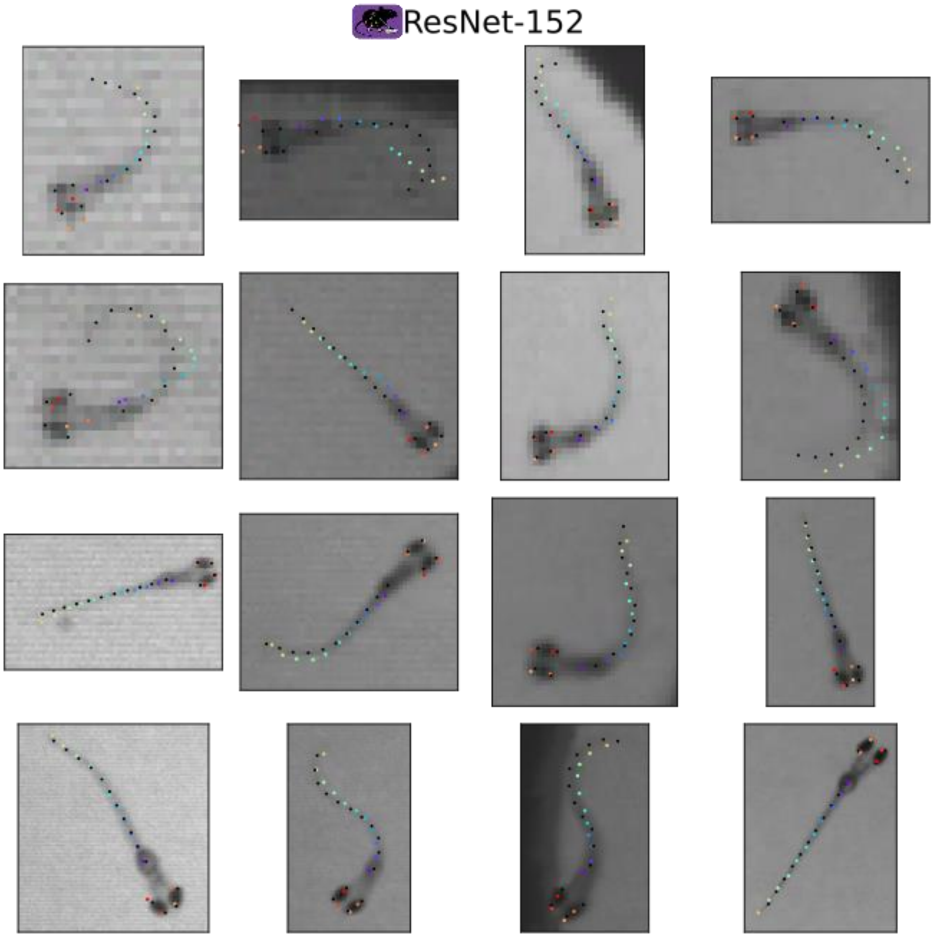
ResNet-152

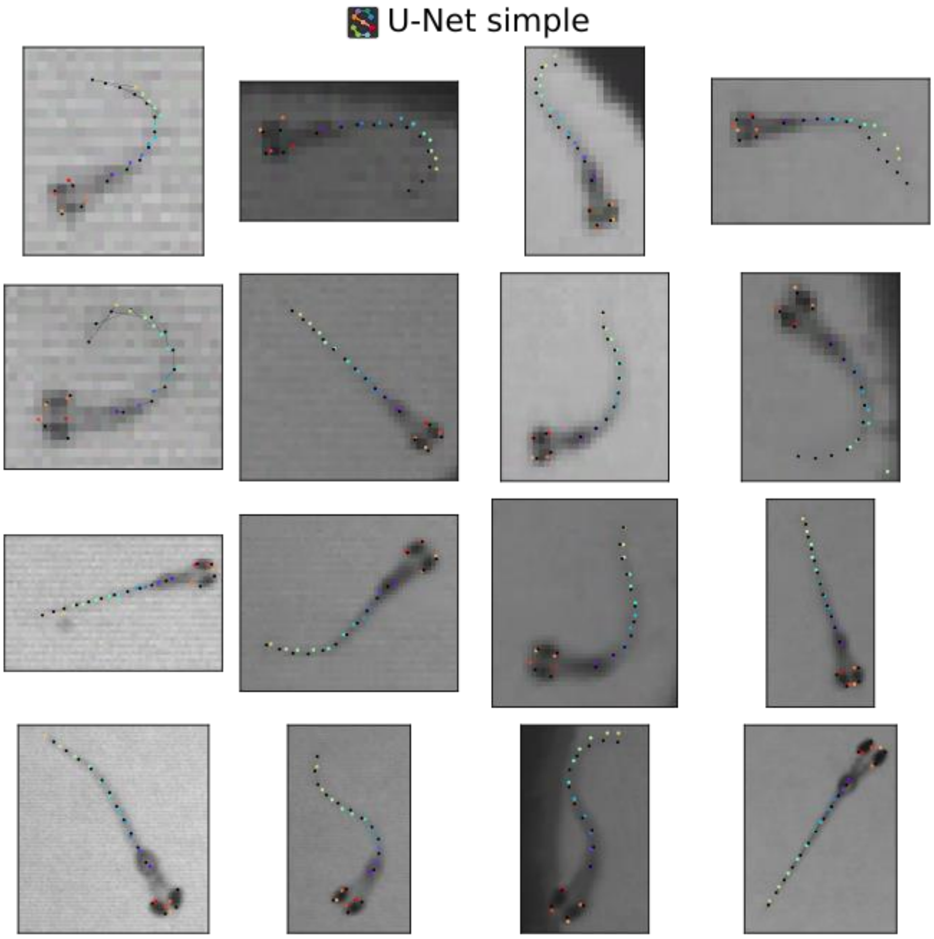


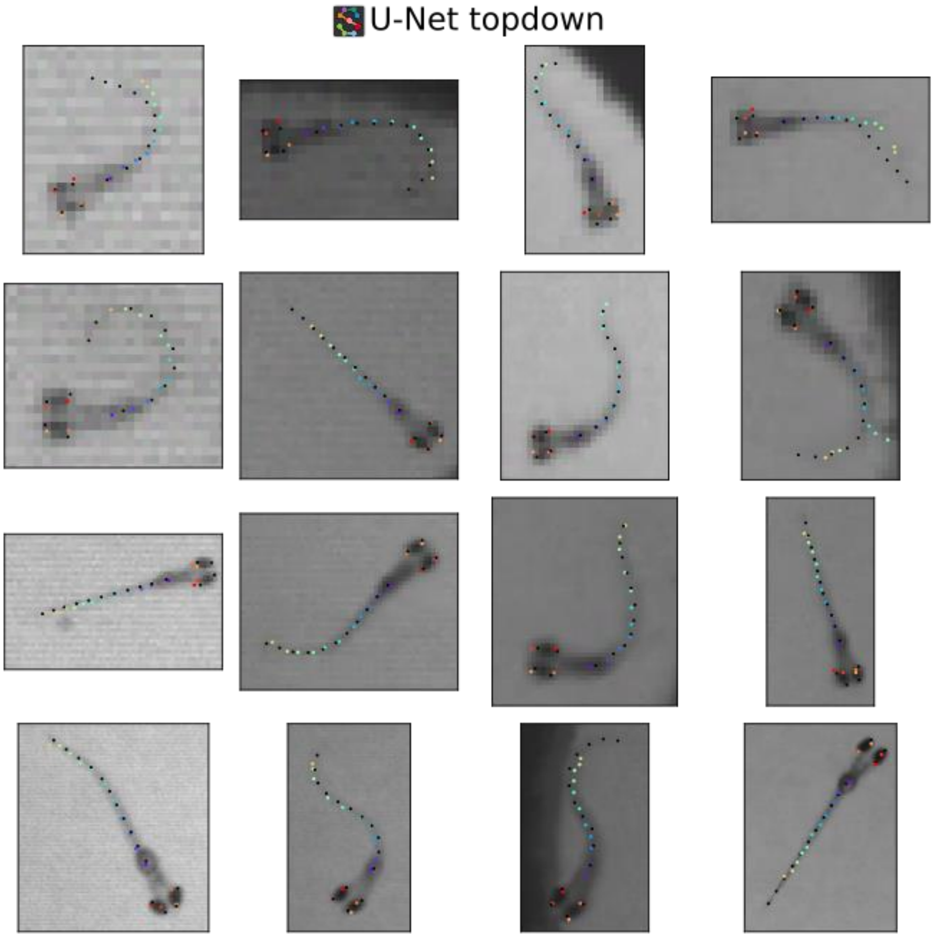


**Figure S2** – Examples of annotated frames (ground truth, shown in the first 4x4 panel) and each model's predictions (DLC Resnet-50, DLC Resnet-152, SLEAP simple, and SLEAP topdown). Columns in the grids represent different camera exposure times (ms) and rows represent different spatial resolutions. For each network panel (4x4 grid), black keypoints represent the ground truth annotations.

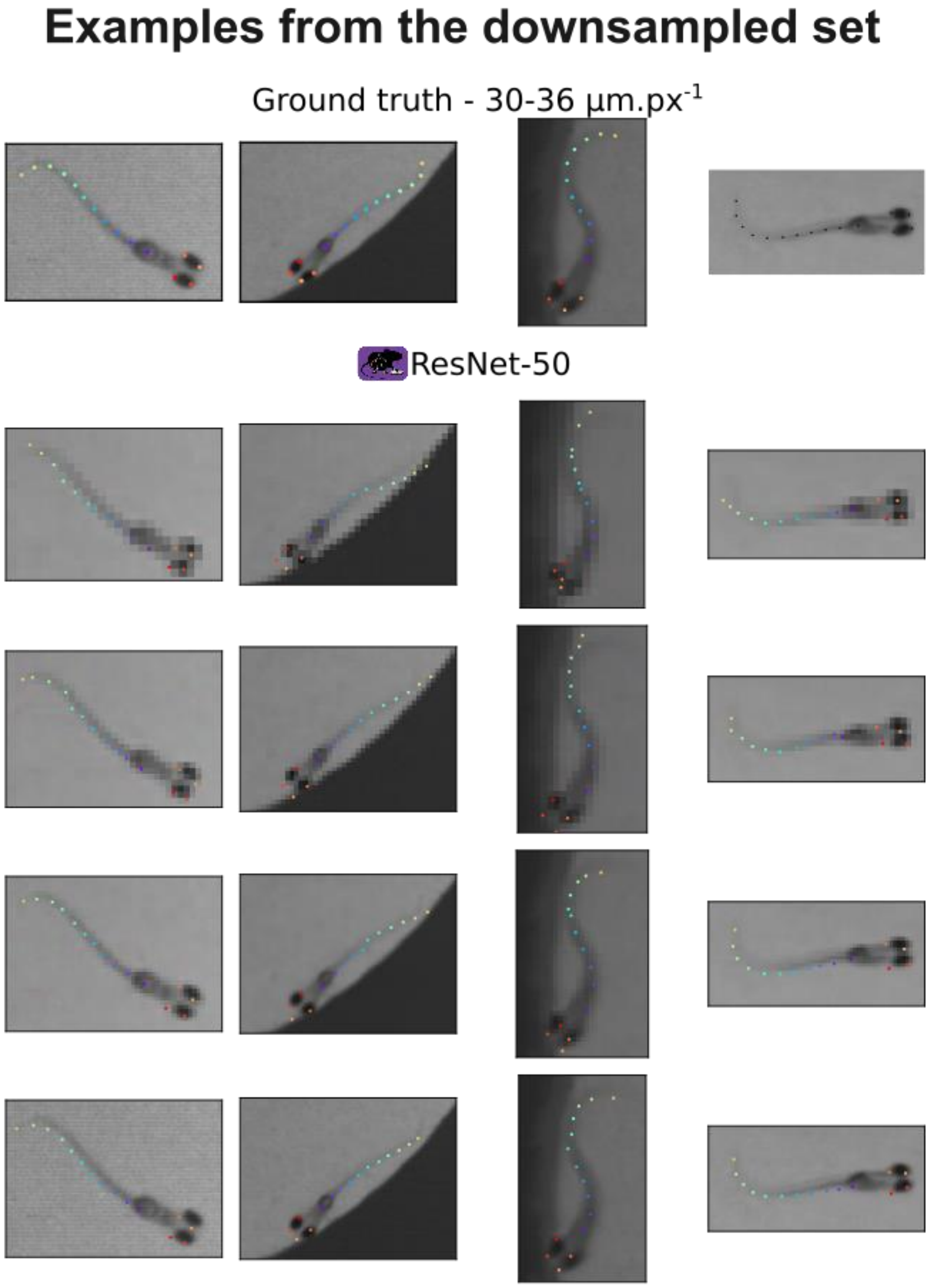

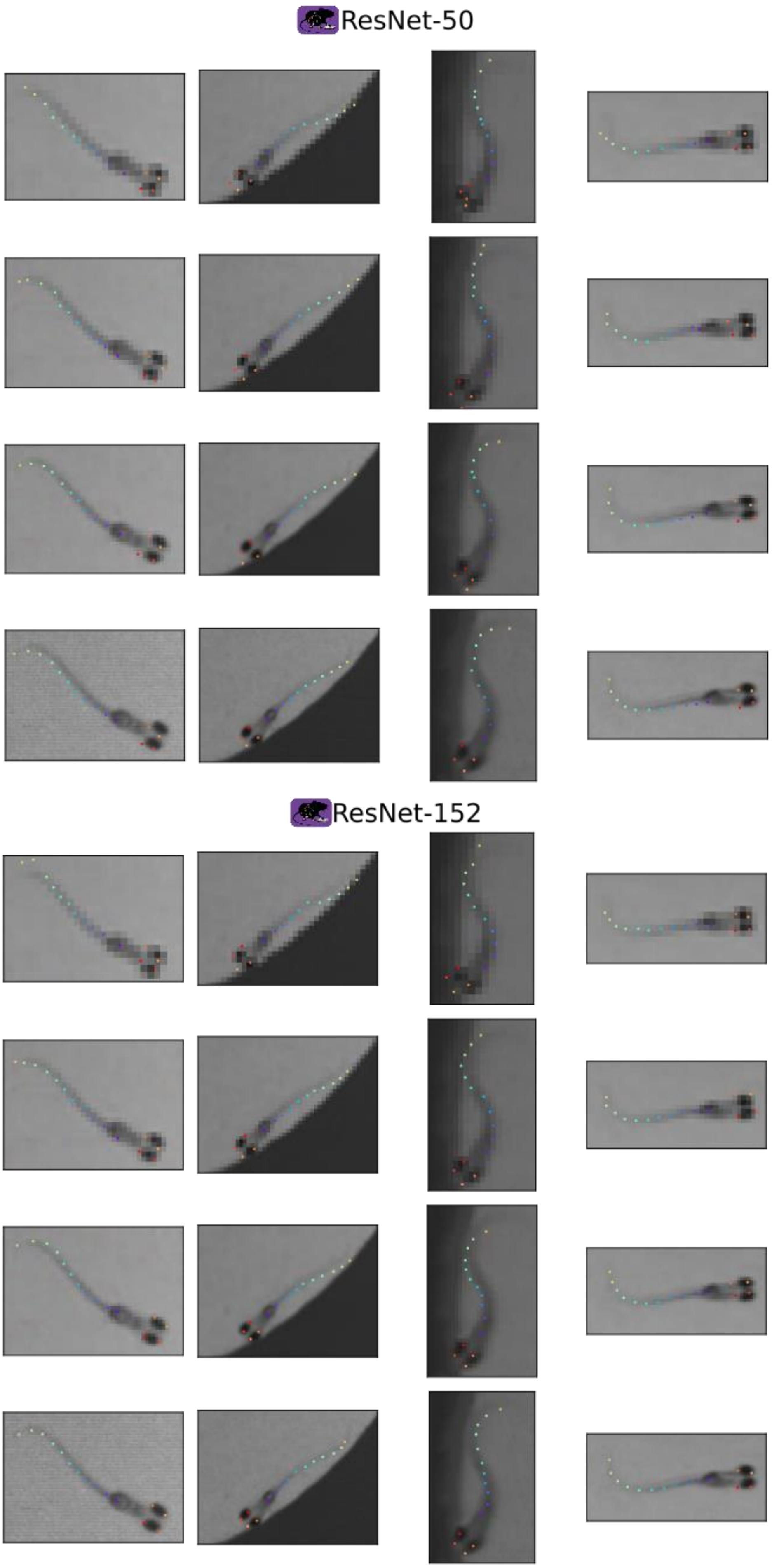
ResNet-50
ResNet-152

U-Net simple

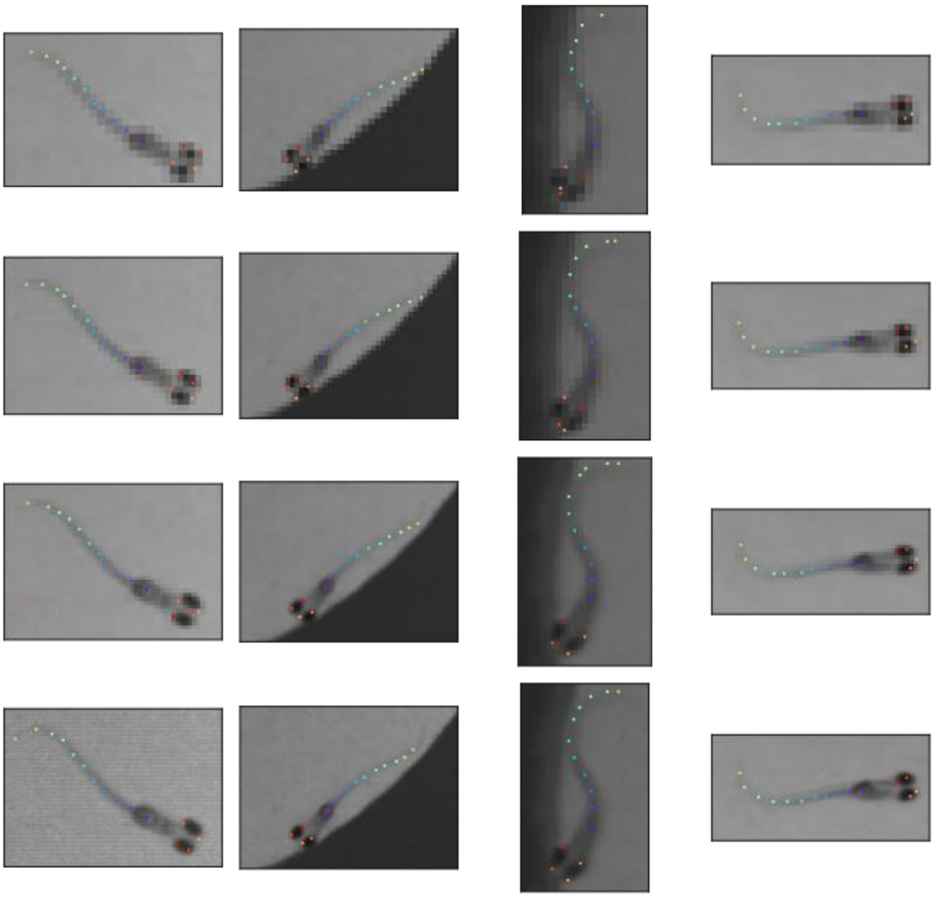

U-Net topdown

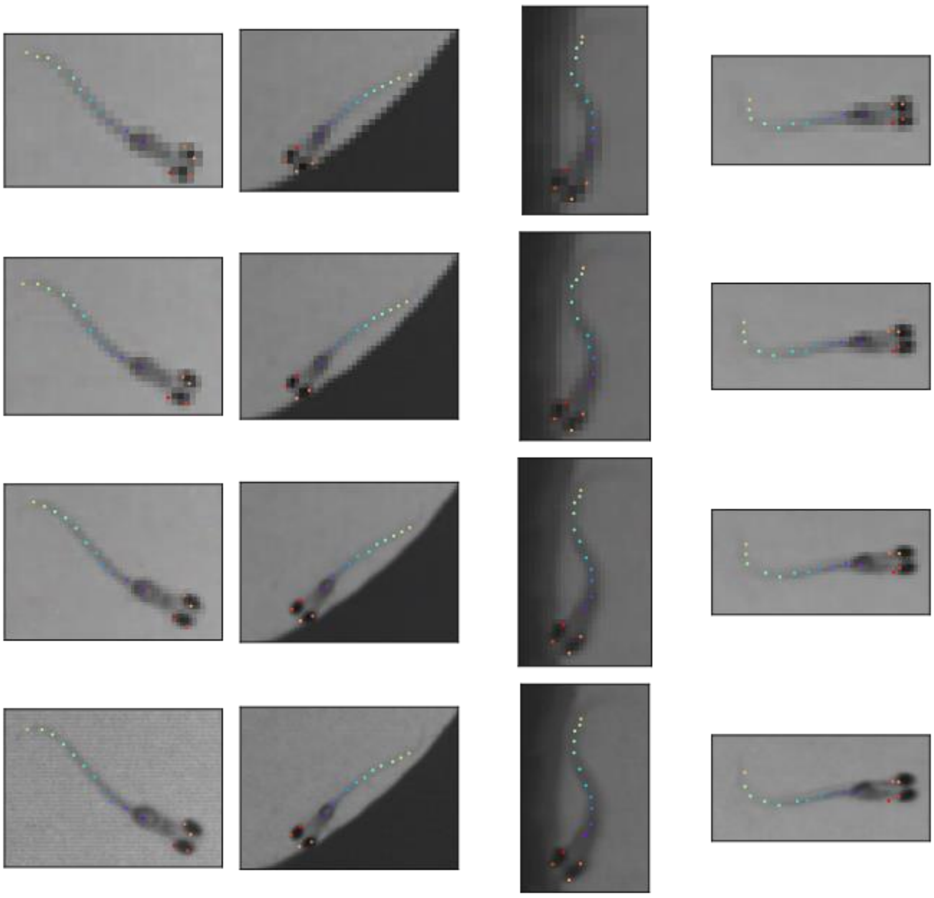

**Figure S3** – Examples of annotated frames and each model's predictions from the dataset built by downsampling the high-resolution frames. There are only 4 ground truth frames, shown first, four panels for each network (DLC Resnet-50, Resnet-152, SLEAP simple, and SLEAP topdown) Columns in the grid represent different camera exposure times (ms) and rows represent different spatial resolutions. For each network panel (4x4 grid), black keypoints represent the ground truth annotations.

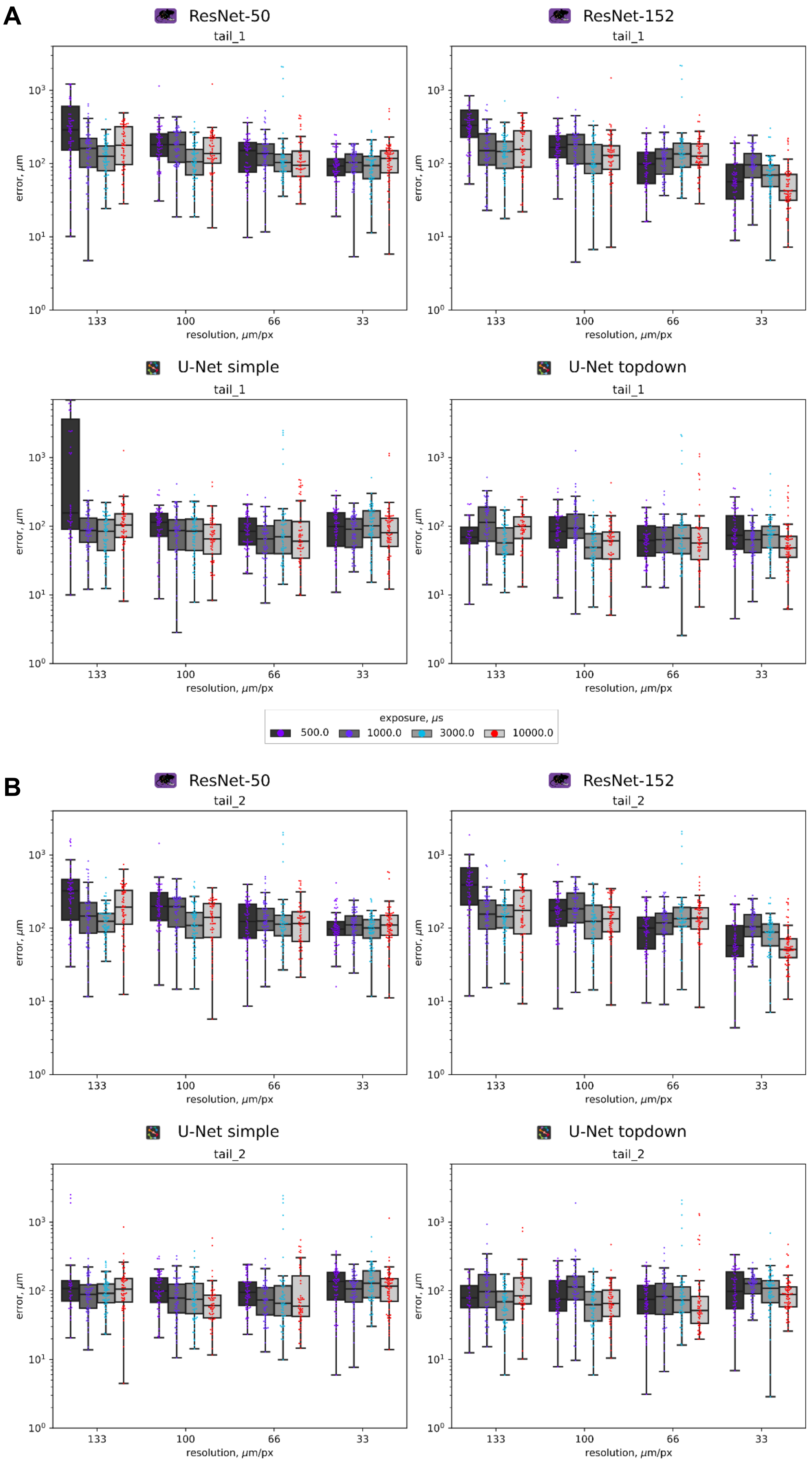


**Figure S4 A-B** – Euclidean distance errors of “tail_1” (**A**) and “tail_2” (**B**) keypoints from each of the four pre-trained networks grouped by spatial resolution using the independent dataset.

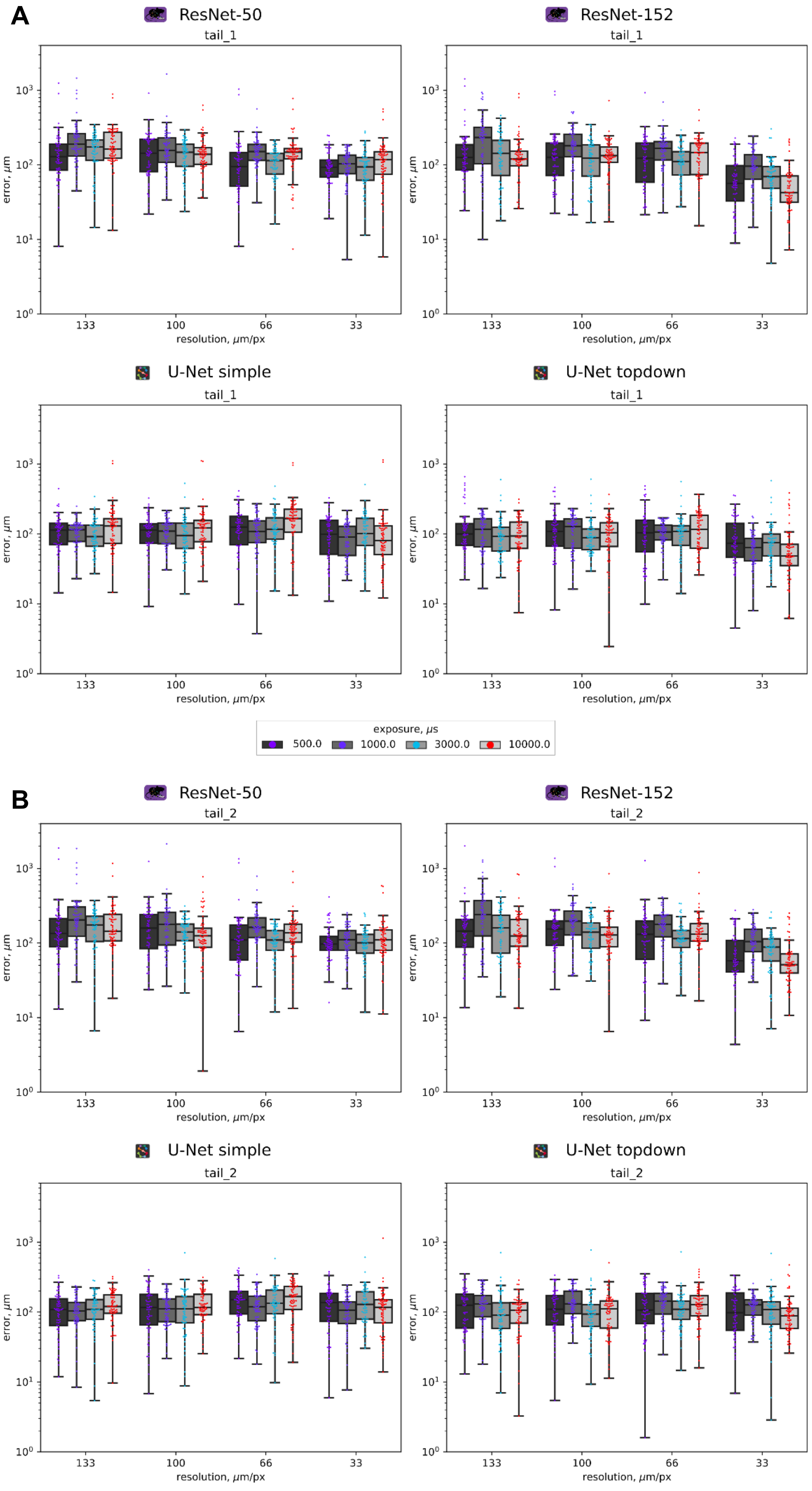


**Figure S5 A-B** – Euclidean distance errors of "tail_1" (**A**) and "tail_2" (**B**) keypoints from each of the four pre-trained networks grouped by spatial resolution using the downsampled dataset.

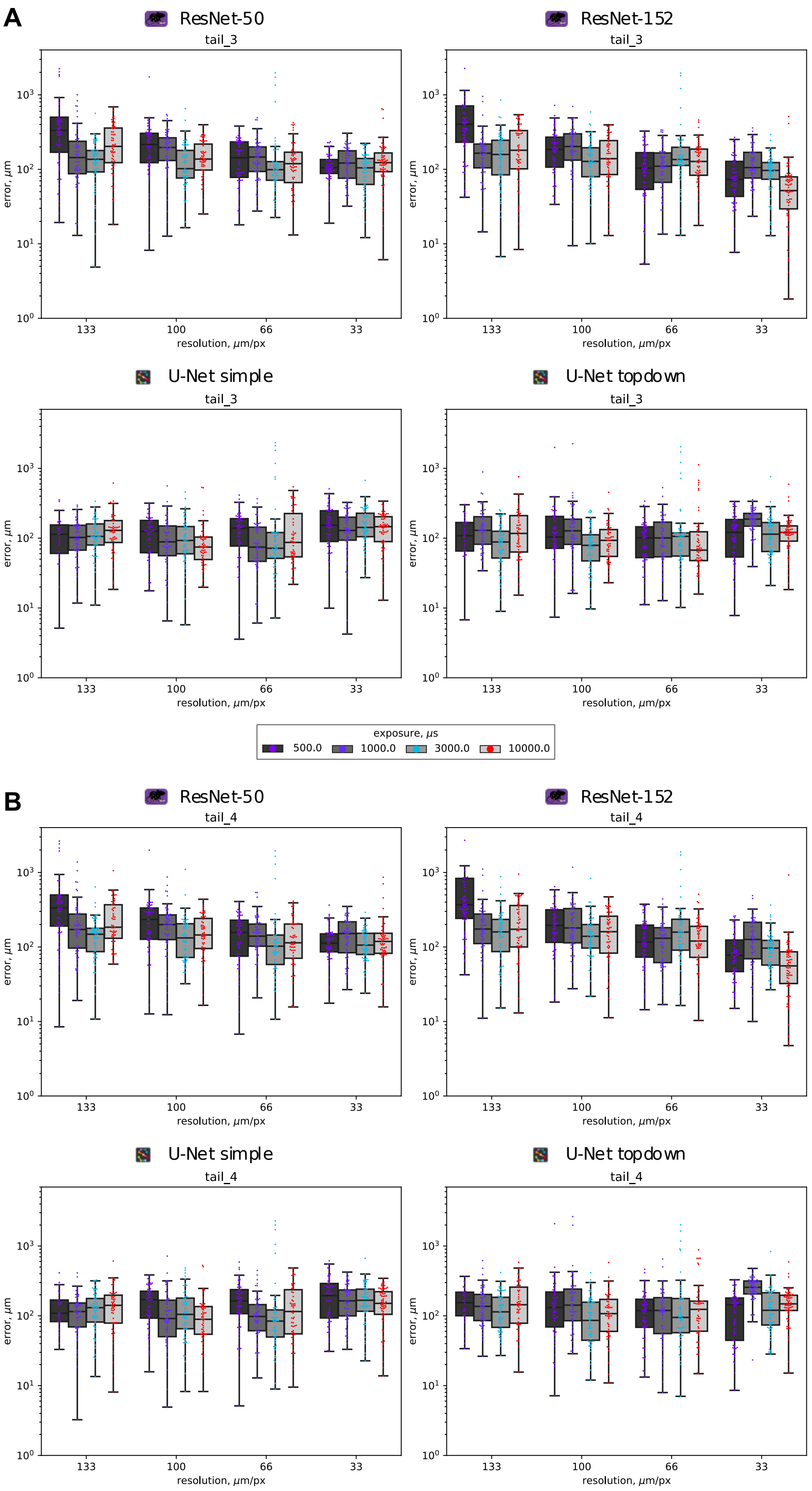


**Figure S6 A-B** – Euclidean distance errors of “tail_3” (**A**) and “tail_4” (**B**) keypoints from each of the four pre-trained networks grouped by spatial resolution using the independent dataset.

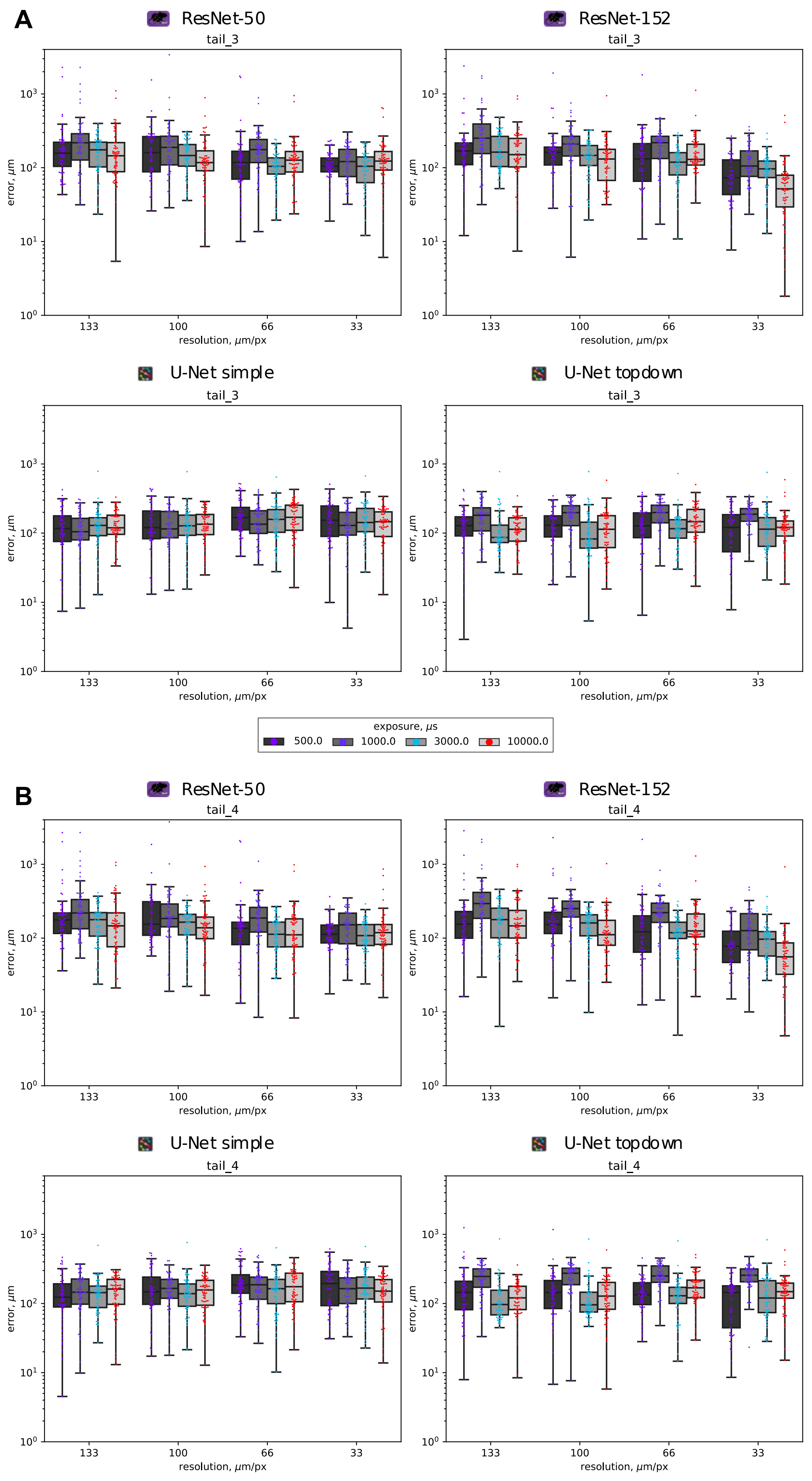


**Figure S7 A-B** – Euclidean distance errors of “tail_3” (**A**) and “tail_4” (**B**) keypoints from each of the four pre-trained networks grouped by spatial resolution using the downsampled dataset.

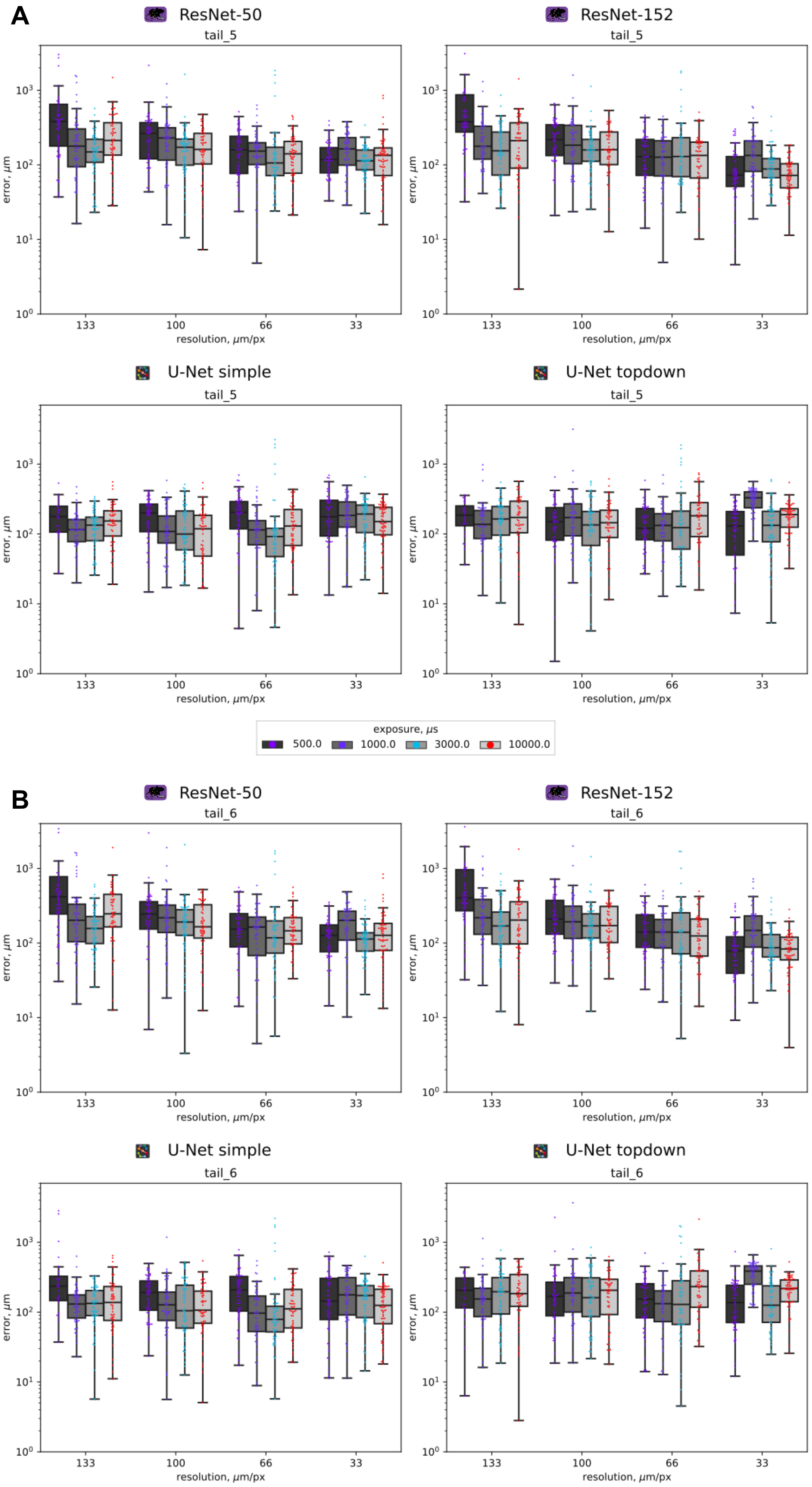


**Figure S8 A-B** – Euclidean distance errors of "tail_5" (**A**) and "tail_6" (**B**) keypoints from each of the four pre-trained networks grouped by spatial resolution using the independent dataset.

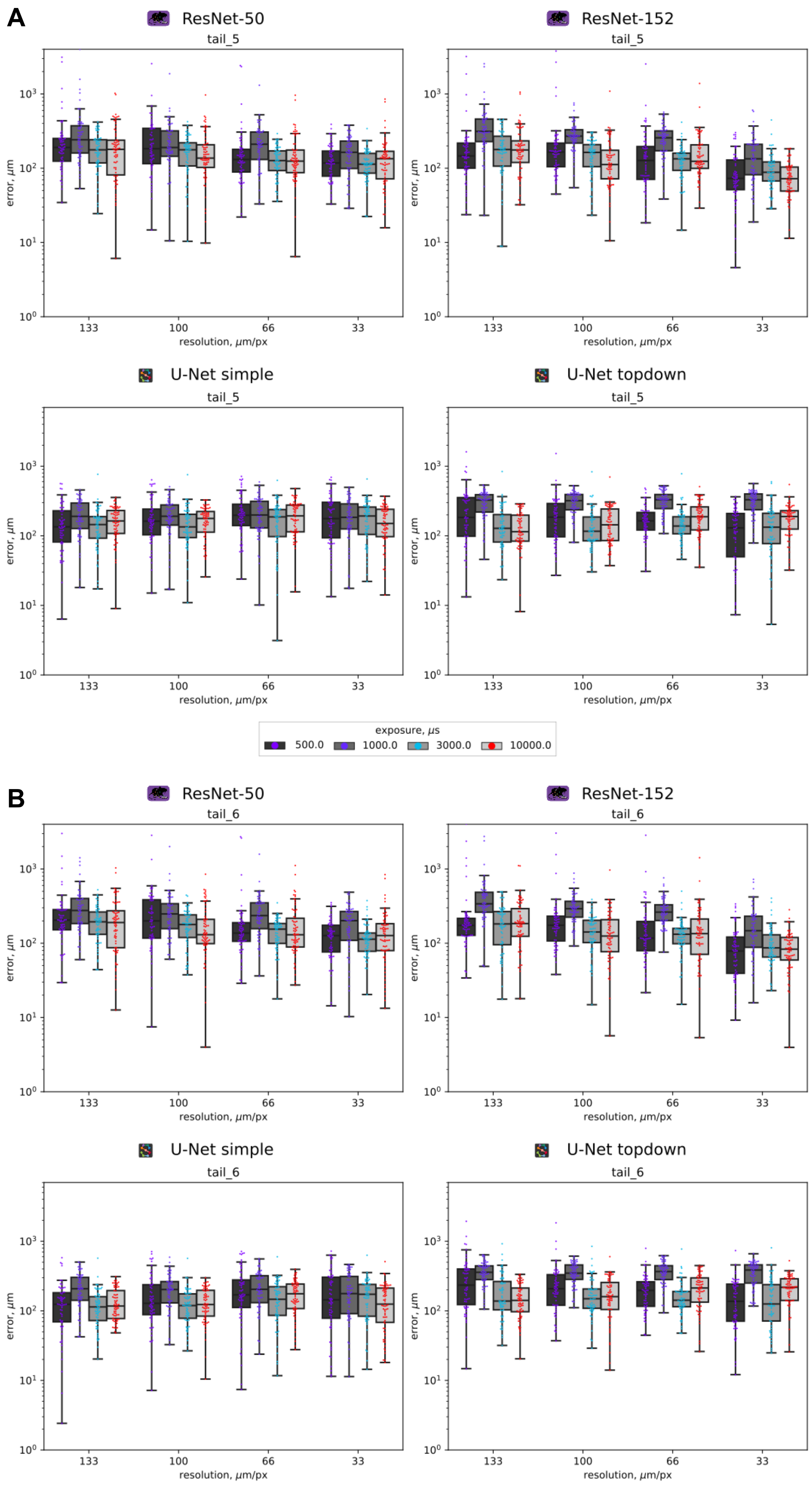


**Figure S9 A-B** – Euclidean distance errors of "tail_5" (**A**) and "tail_6" (**B**) keypoints from each of the four pre-trained networks grouped by spatial resolution using the downsampled dataset.

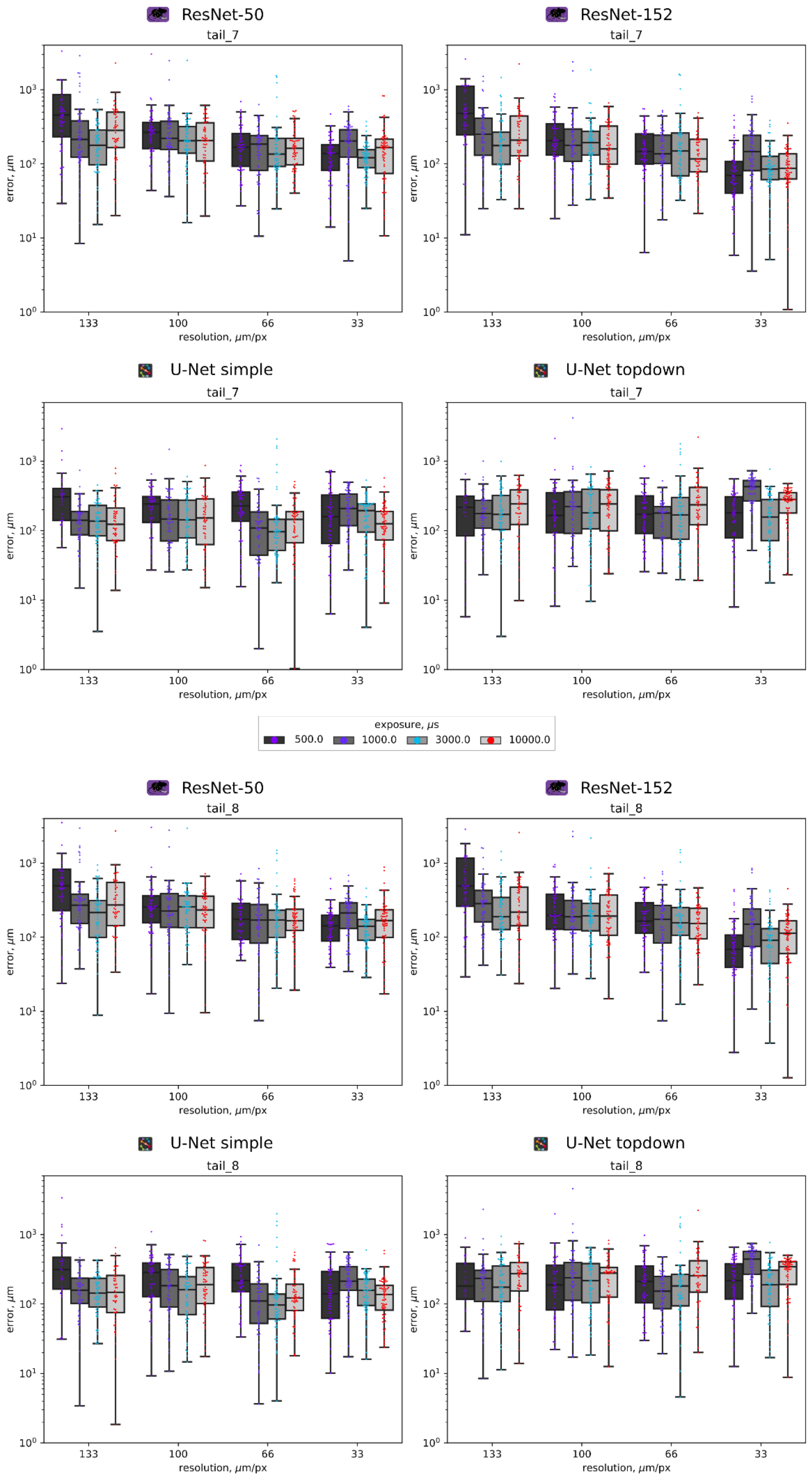


**Figure S10 A-B** – Euclidean distance errors of "tail_7" (**A**) and "tail_8" (**B**) keypoints from each of the four pre-trained networks grouped by spatial resolution using the independent dataset.

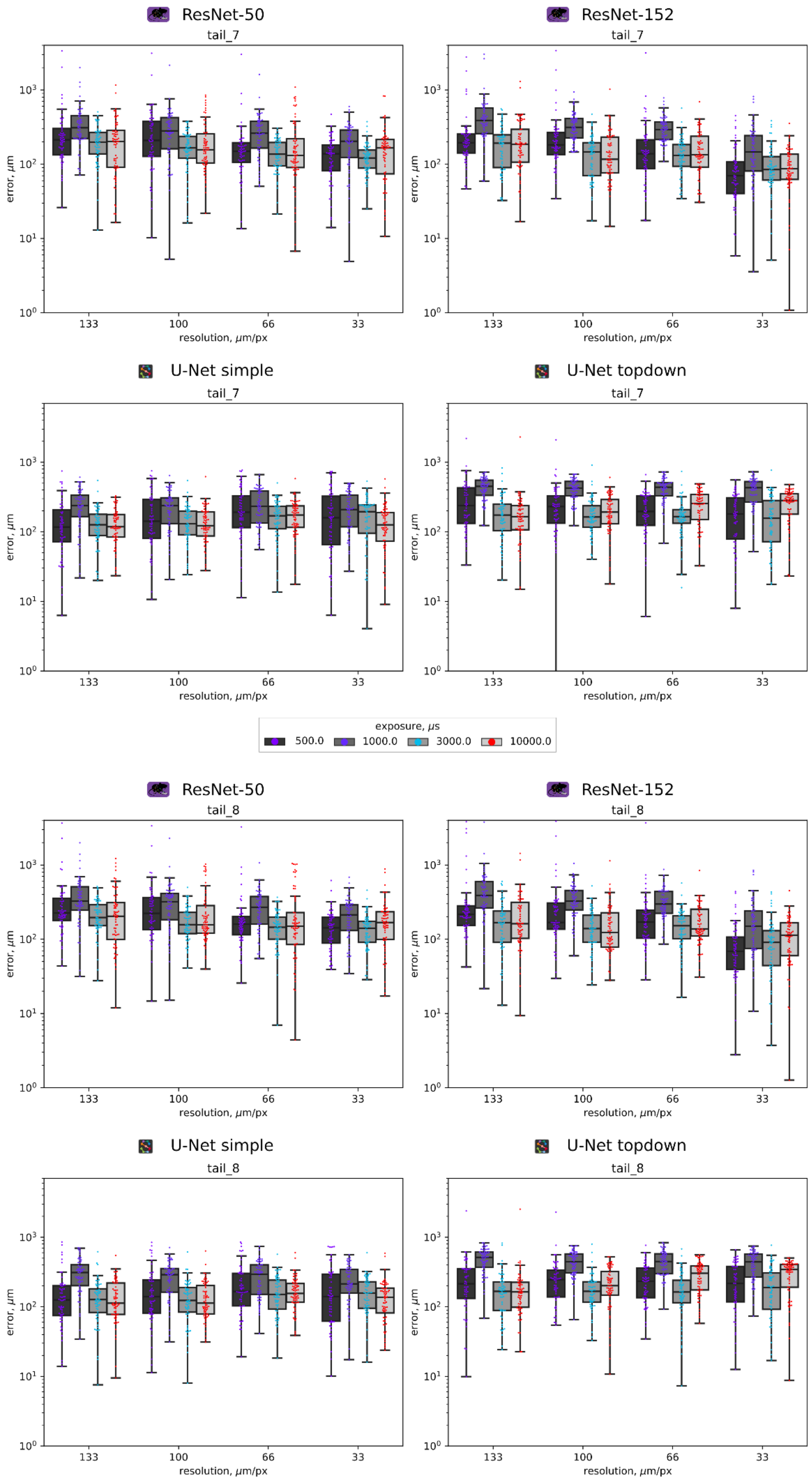


**Figure S11 A-B** – Euclidean distance errors of "tail_7" (**A**) and "tail_8" (**B**) keypoints from each of the four pre-trained networks grouped by spatial resolution using the downsampled dataset.

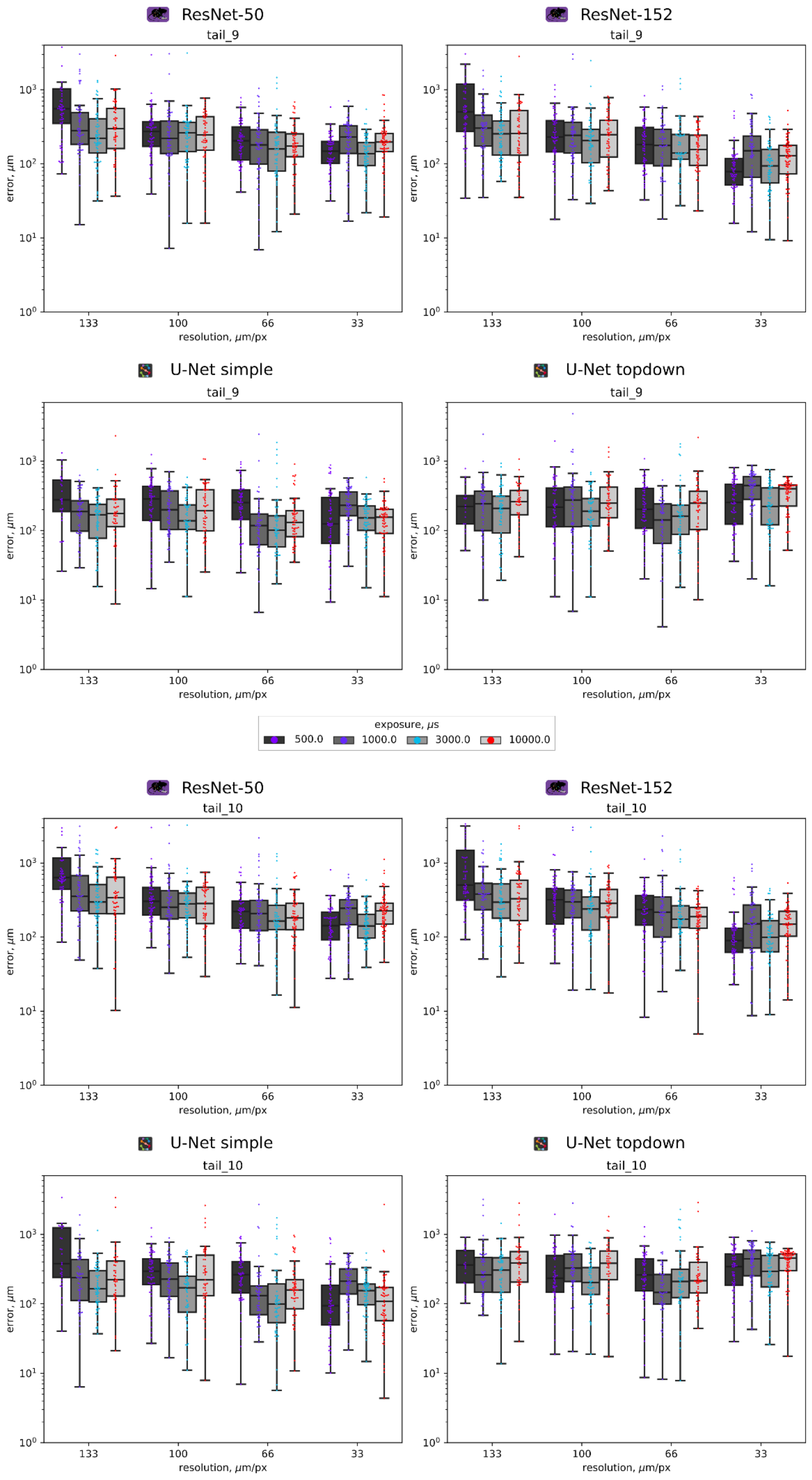


**Figure S10 A-B** – Euclidean distance errors of "tail_9" (**A**) and "tail_10" (**B**) keypoints from each of the four pre-trained networks grouped by spatial resolution using the independent dataset.

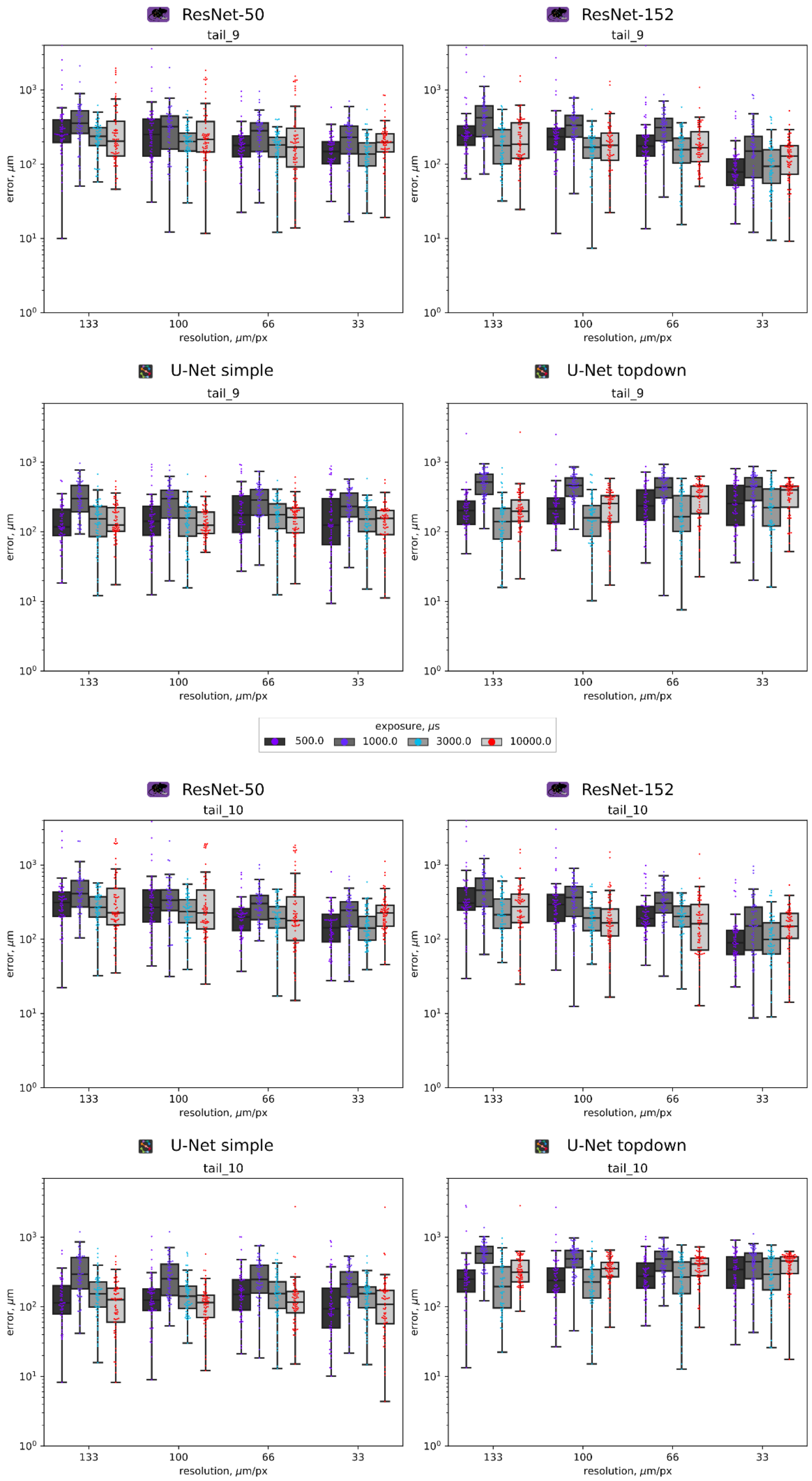


**Figure S11 A-B** – Euclidean distance errors of “tail_9” (**A**) and “tail_10” (**B**) keypoints from each of the four pre-trained networks grouped by spatial resolution using the downsampled dataset.

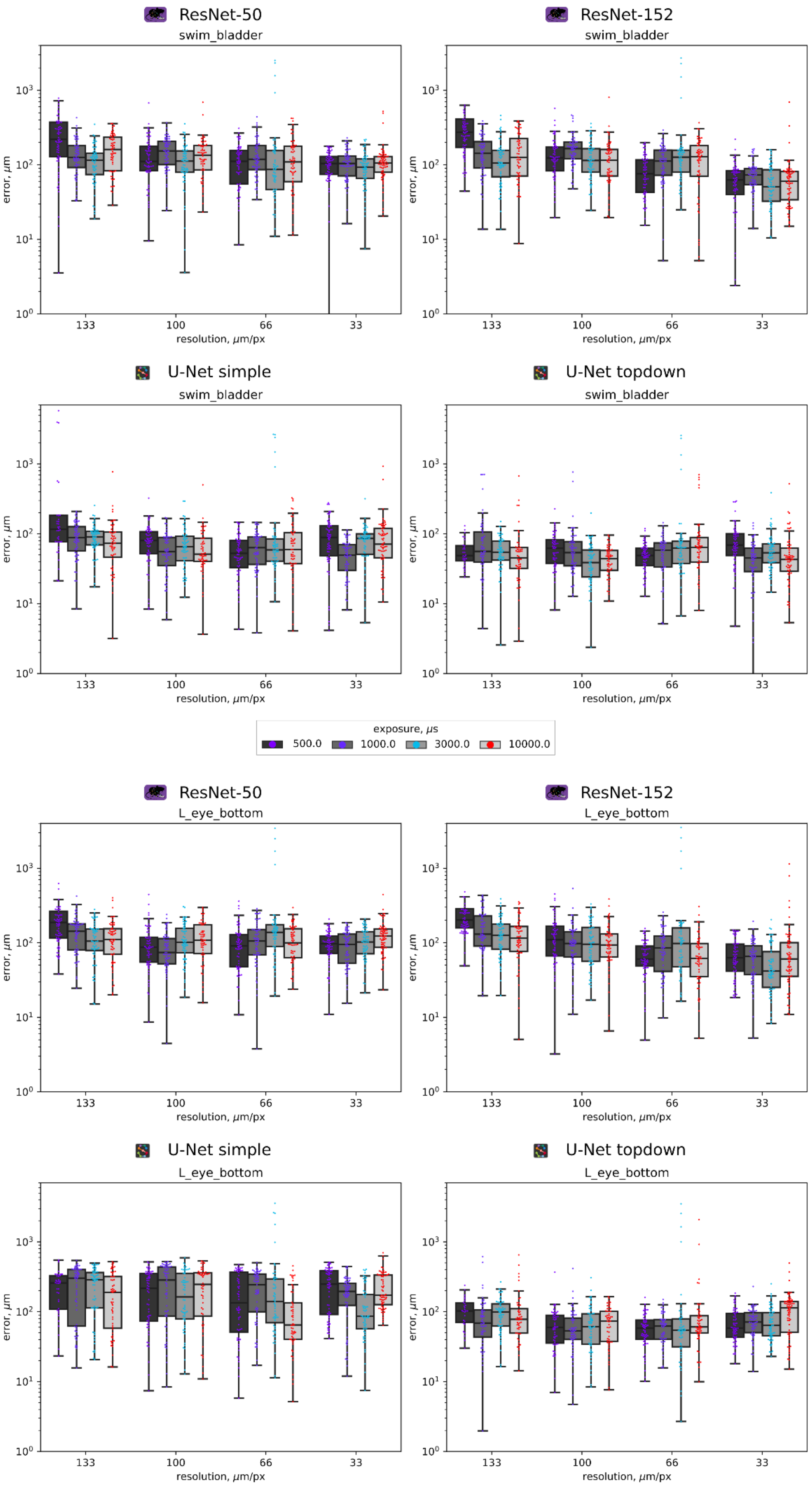


**Figure S12 A-B** – Euclidean distance errors of "swim_bladder" (**A**) and "Left(L)_eye_bottom" (**B**) keypoints from each of the four pre-trained networks grouped by spatial resolution using the independent dataset.

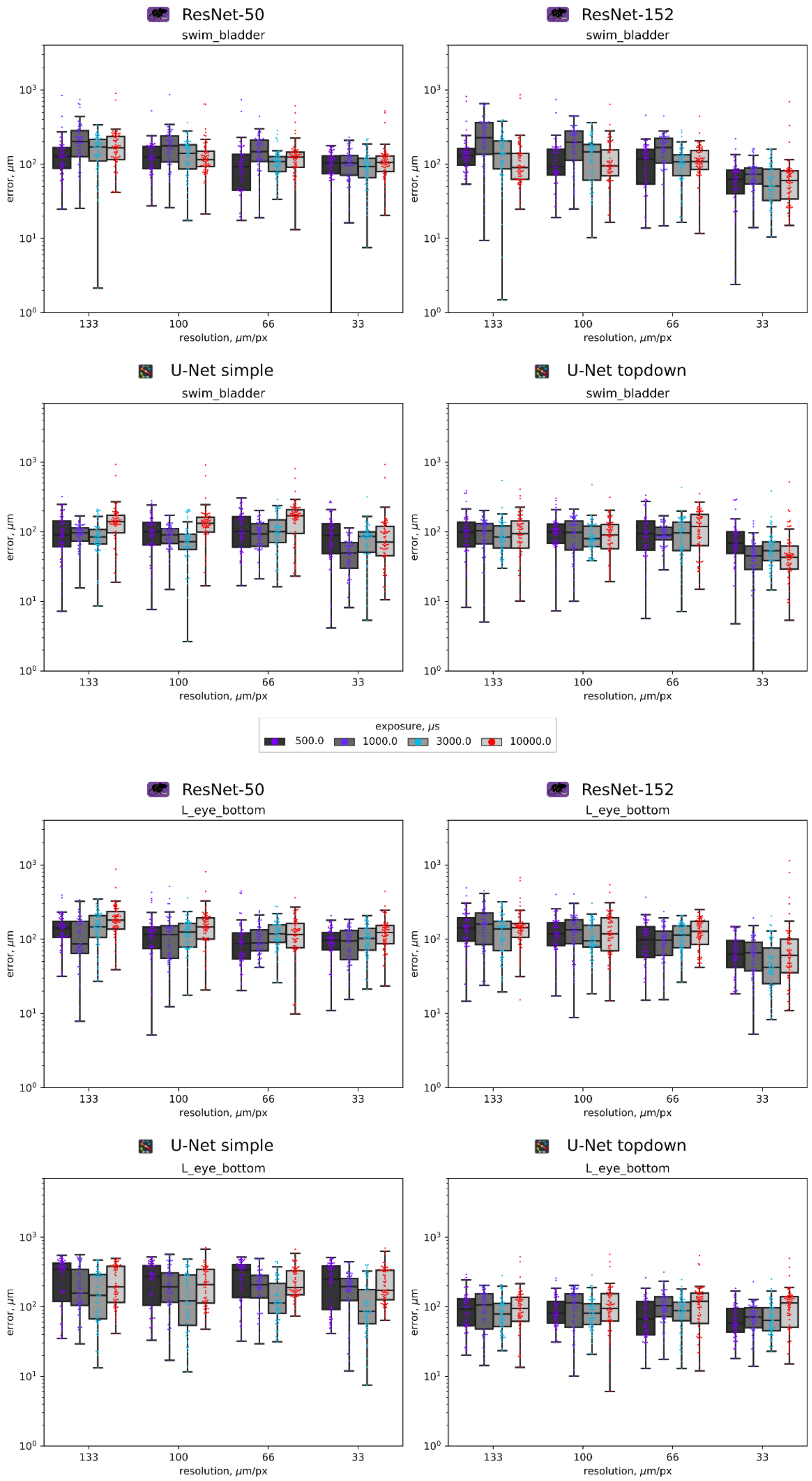


**Figure S13 A-B** – Euclidean distance errors of "swim_bladder" (**A**) and "Left(L)_eye_bottom" (**B**) keypoints from each of the four pre-trained networks grouped by spatial resolution using the downsampled dataset.

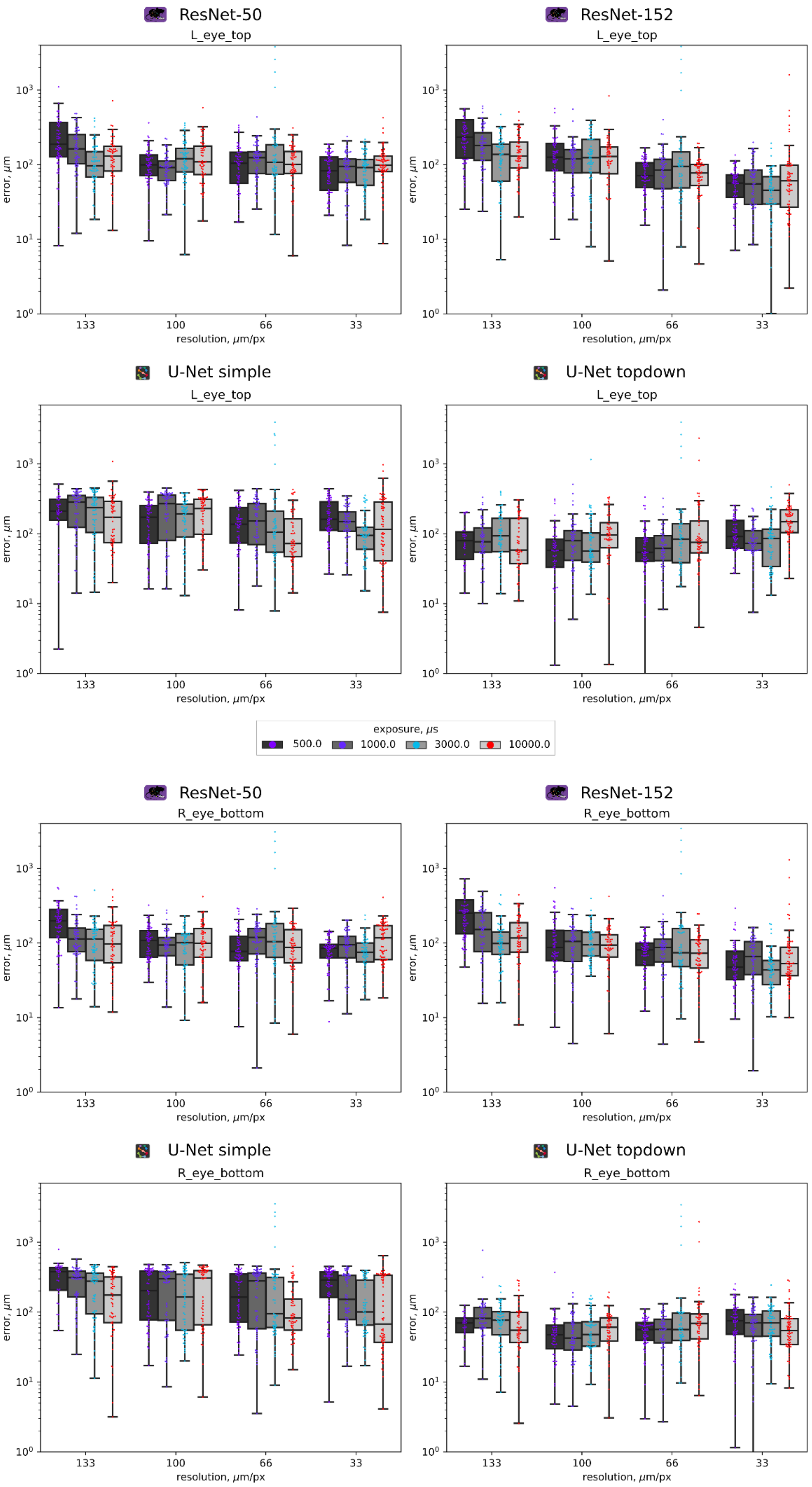


**Figure S14 A-B** – Euclidean distance errors of "Left(L)_eye_top" (**A**) and "Right(R)_eye_bottom" (**B**) keypoints from each of the four pre-trained networks grouped by spatial resolution using the independent dataset.

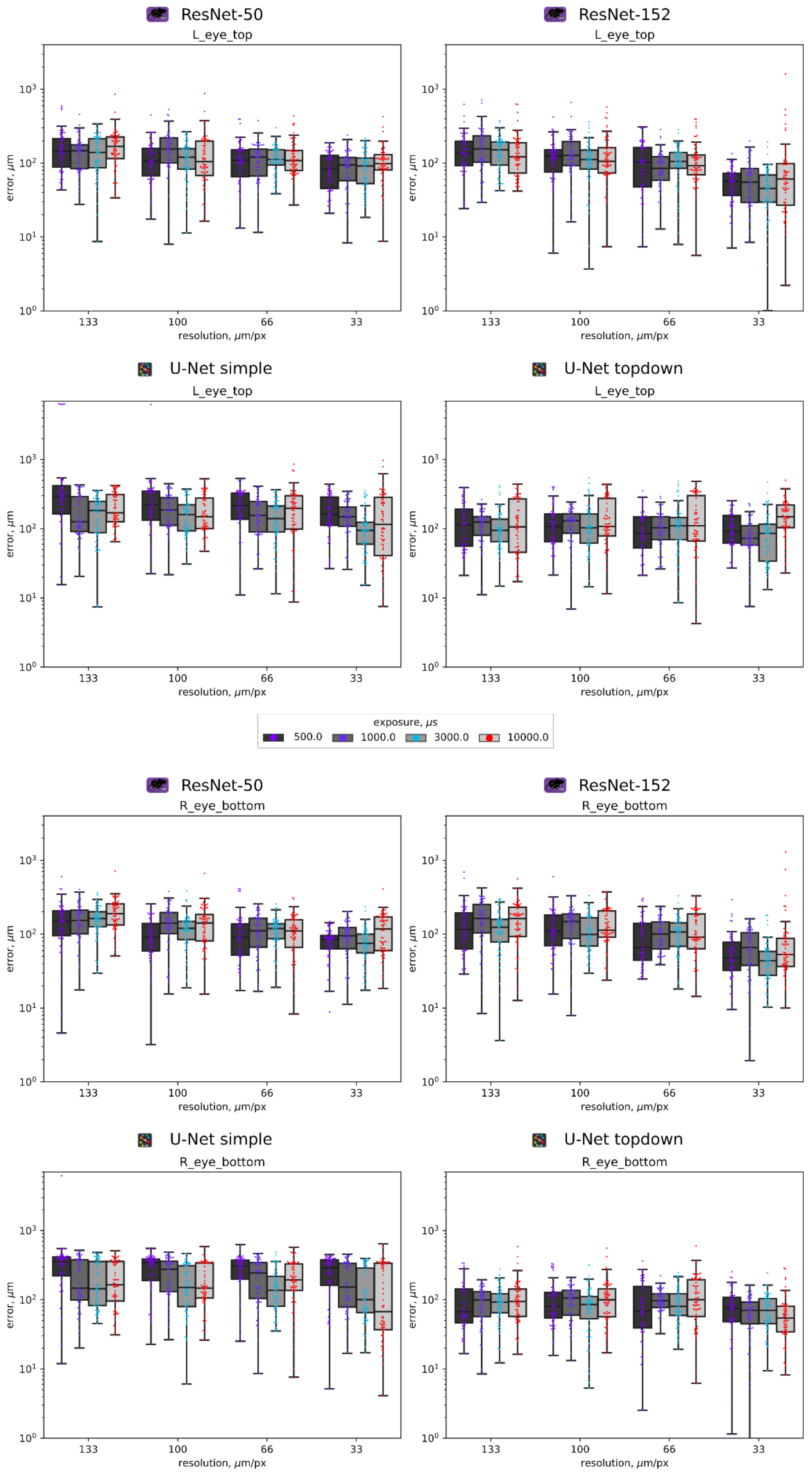


**Figure S15 A-B** – Euclidean distance errors of “Left(L)_eye_top” (**A**) and “Right(R)_eye_bottom” (**B**) keypoints from each of the four pre-trained networks grouped by spatial resolution using the independent dataset.

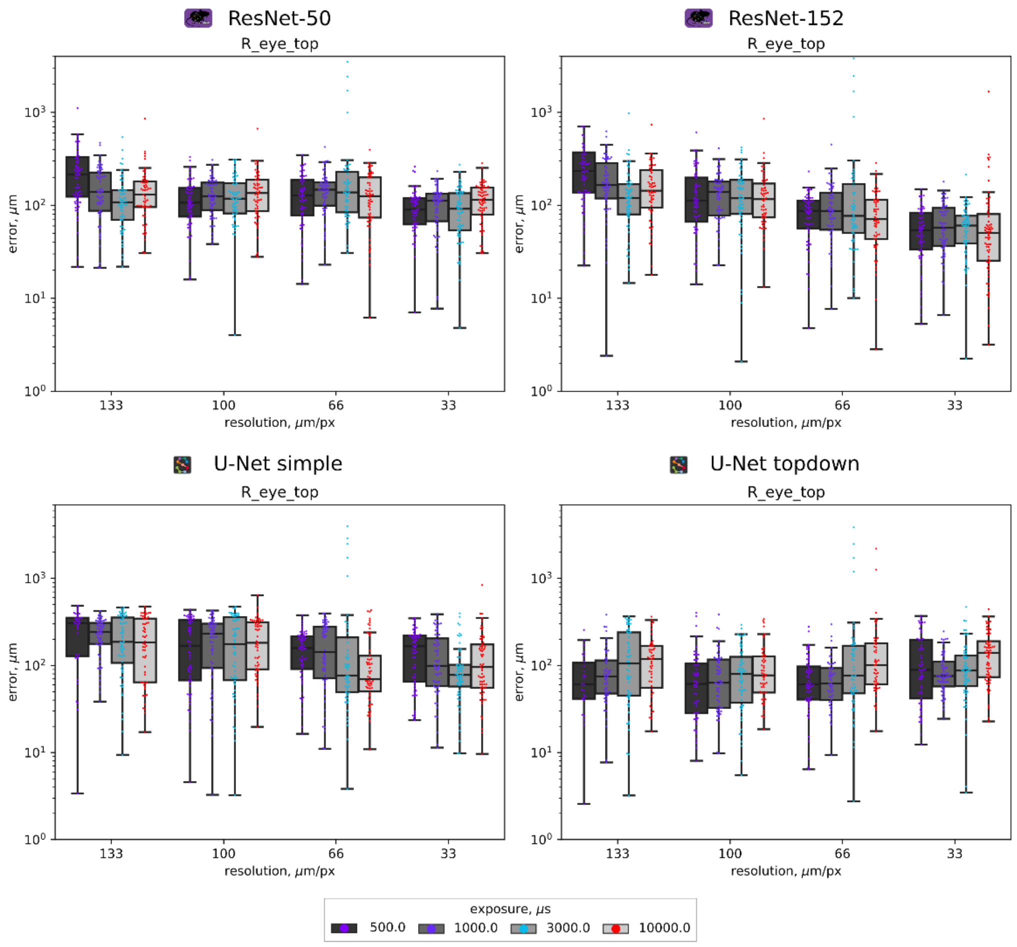


**Figure S16 A-B** – Euclidean distance errors of "Left(L)_eye_top" (**A**) and "Right(R)_eye_bottom" (**B**) keypoints from each of the four pre-trained networks grouped by spatial resolution using the independent dataset.

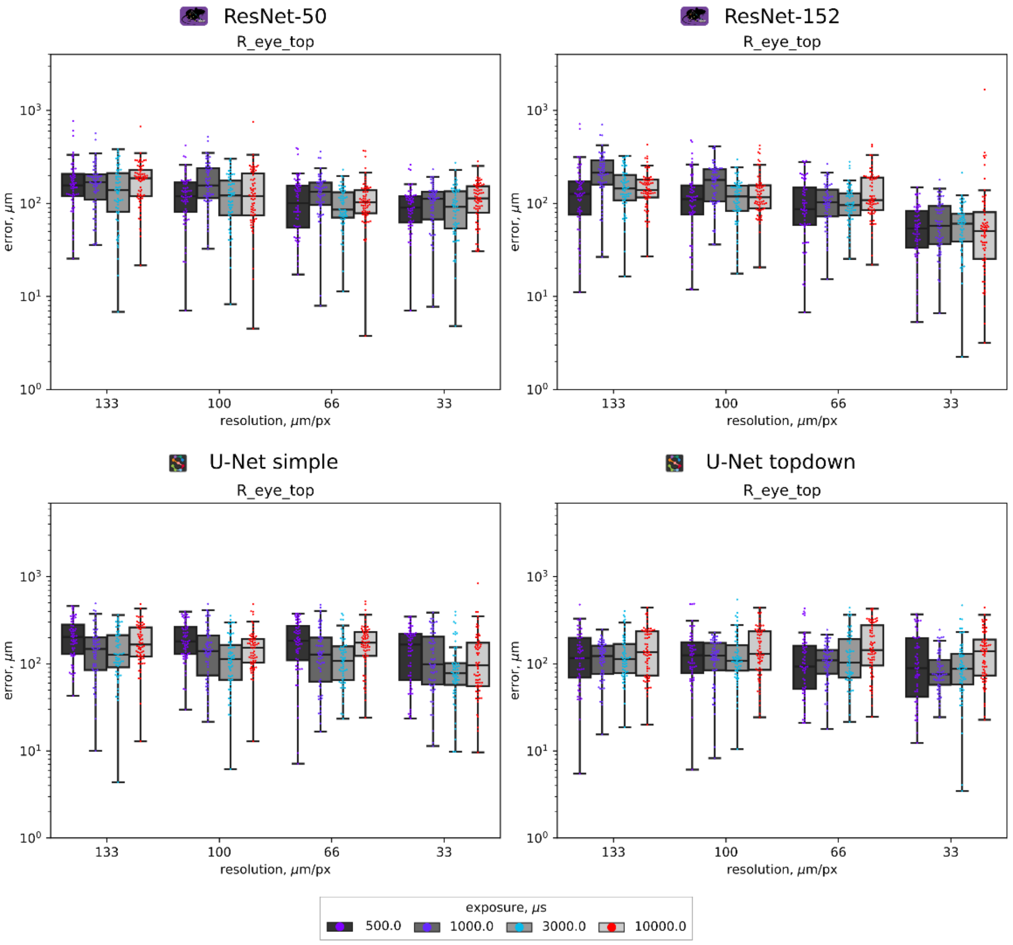


**Figure S17 A-B** – Euclidean distance errors of "Left(L)_eye_top" (**A**) and "Right(R)_eye_bottom" (**B**) keypoints from each of the four pre-trained networks grouped by spatial resolution using the independent dataset.

**Table ST1 A-D** – p-values for each pairwise comparisons of the mean Euclidean distance errors, which include all keypoints, per resolution and camera exposure groups for both the independent (**A** and **B**) and downsampled (**C** and **D**) benchmarking datasets. At the resolution tables, the numbers 1-4 represent 130-136, 97-103, 63-69 and 30-36 µm/px resolutions, respectively. At the exposure tables, the numbers 1-4 represent 0.5, 1, 3 and 10 ms exposure times, respectively. Yellow ($0.01 \leq p < 0.05$), orange ($0.002 \leq p < 0.01$) and light red ($p < 0.002$) values depict significant differences between those pairs of data (see colormap).

**A**

**Resolution**

| Independent dataset | 1.0_DLC_Resnet-152 | 1.0_DLC_Resnet-50 | 1.0_SLEAP_UNet_simple | 1.0_SLEAP_UNet_topdown | 2.0_DLC_Resnet-152 | 2.0_DLC_Resnet-50 | 2.0_SLEAP_UNet_simple | 2.0_SLEAP_UNet_topdown | 3.0_DLC_Resnet-152 | 3.0_DLC_Resnet-50 | 3.0_SLEAP_UNet_simple | 3.0_SLEAP_UNet_topdown | 4.0_DLC_Resnet-152 | 4.0_DLC_Resnet-50 | 4.0_SLEAP_UNet_simple | 4.0_SLEAP_UNet_topdown |
|---|---|---|---|---|---|---|---|---|---|---|---|---|---|---|---|---|
| 1.0_DLC_Resnet-152 | 1 | | | | | | | | | | | | | | | |
| 1.0_DLC_Resnet-50 | 1 | 1 | | | | | | | | | | | | | | |
| 1.0_SLEAP_UNet_simple | 0.00199178 | 0.02202577 | 1 | | | | | | | | | | | | | |
| 1.0_SLEAP_UNet_topdown | 3.22E-11 | 1.56E-09 | 0.20810305 | 1 | | | | | | | | | | | | |
| 2.0_DLC_Resnet-152 | 0.00208977 | 0.02349601 | 1 | 0.15933066 | 1 | | | | | | | | | | | |
| 2.0_DLC_Resnet-50 | 0.09342537 | 0.6344673 | 1 | 0.00524828 | 1 | 1 | | | | | | | | | | |
| 2.0_SLEAP_UNet_simple | 7.35E-06 | 0.00015286 | 1 | 1 | 1 | 1 | 1 | | | | | | | | | |
| 2.0_SLEAP_UNet_topdown | 1.75E-16 | 2.24E-14 | 0.00117785 | 1 | 0.00075961 | 6.91E-06 | 0.07381862 | 1 | | | | | | | | |
| 3.0_DLC_Resnet-152 | 5.38E-21 | 1.35E-18 | 6.09E-06 | 1 | 3.35E-06 | 1.14E-08 | 0.00102347 | 1 | 1 | | | | | | | |
| 3.0_DLC_Resnet-50 | 1.53E-13 | 1.32E-11 | 0.03940298 | 1 | 0.02807655 | 0.00050621 | 1 | 1 | 1 | 1 | | | | | | |
| 3.0_SLEAP_UNet_simple | 3.68E-18 | 6.26E-16 | 0.00019876 | 1 | 0.0001209 | 7.48E-07 | 0.01815072 | 1 | 1 | 1 | 1 | | | | | |
| 3.0_SLEAP_UNet_topdown | 5.52E-25 | 2.04E-22 | 1.94E-08 | 0.28941728 | 9.32E-09 | 1.52E-11 | 7.05E-06 | 1 | 1 | 0.47701167 | 1 | 1 | | | | |
| 4.0_DLC_Resnet-152 | 9.02E-70 | 2.48E-65 | 6.08E-38 | 3.32E-19 | 5.07E-39 | 1.08E-44 | 9.94E-33 | 5.97E-16 | 1.08E-12 | 5.98E-20 | 4.94E-15 | 6.94E-09 | 1 | | | |
| 4.0_DLC_Resnet-50 | 3.69E-27 | 2.16E-24 | 2.82E-09 | 0.16555504 | 1.23E-09 | 1.21E-12 | 1.59E-06 | 1 | 1 | 0.2719017 | 1 | 1 | 2.50E-09 | 1 | | |
| 4.0_SLEAP_UNet_simple | 9.36E-10 | 4.49E-08 | 1 | 1 | 1 | 0.07956626 | 1 | 1 | 0.19125523 | 1 | 1 | 0.00379791 | 1.82E-26 | 0.00140522 | 1 | |
| 4.0_SLEAP_UNet_topdown | 0.00034696 | 0.00493398 | 1 | 0.41615184 | 1 | 1 | 1 | 0.00279484 | 1.57E-05 | 0.08581036 | 0.00048975 | 5.19E-08 | 7.19E-38 | 7.61E-09 | 1 | 1 |

**B**

**Exposure**

| Independent dataset | 1.0_DLC_Resnet-152 | 1.0_DLC_Resnet-50 | 1.0_SLEAP_UNet_simple | 1.0_SLEAP_UNet_topdown | 2.0_DLC_Resnet-152 | 2.0_DLC_Resnet-50 | 2.0_SLEAP_UNet_simple | 2.0_SLEAP_UNet_topdown | 3.0_DLC_Resnet-152 | 3.0_DLC_Resnet-50 | 3.0_SLEAP_UNet_simple | 3.0_SLEAP_UNet_topdown | 4.0_DLC_Resnet-152 | 4.0_DLC_Resnet-50 | 4.0_SLEAP_UNet_simple | 4.0_SLEAP_UNet_topdown |
|---|---|---|---|---|---|---|---|---|---|---|---|---|---|---|---|---|
| 1.0_DLC_Resnet-152 | 1 | | | | | | | | | | | | | | | |
| 1.0_DLC_Resnet-50 | 1 | 1 | | | | | | | | | | | | | | |
| 1.0_SLEAP_UNet_simple | 0.0738272 | 1 | 1 | | | | | | | | | | | | | |
| 1.0_SLEAP_UNet_topdown | 0.1399741 | 1.14E-05 | 1.65E-08 | 1 | | | | | | | | | | | | |
| 2.0_DLC_Resnet-152 | 1 | 1 | 0.05337564 | 0.23642386 | 1 | | | | | | | | | | | |
| 2.0_DLC_Resnet-50 | 1 | 1 | 1 | 0.00019067 | 1 | 1 | | | | | | | | | | |
| 2.0_SLEAP_UNet_simple | 1 | 1 | 0.15019941 | 0.09261134 | 1 | 1 | 1 | | | | | | | | | |
| 2.0_SLEAP_UNet_topdown | 1 | 1 | 0.11538047 | 0.15871788 | 1 | 1 | 1 | 1 | | | | | | | | |
| 3.0_DLC_Resnet-152 | 0.08218015 | 2.89E-06 | 2.50E-09 | 1 | 0.14681771 | 6.38E-05 | 0.05316097 | 0.09663406 | 1 | | | | | | | |
| 3.0_DLC_Resnet-50 | 1 | 0.0004731 | 1.05E-06 | 1 | 1 | 0.00590216 | 1 | 1 | 1 | 1 | | | | | | |
| 3.0_SLEAP_UNet_simple | 1 | 0.00332626 | 1.12E-05 | 1 | 1 | 0.03256969 | 1 | 1 | 1 | 1 | 1 | | | | | |
| 3.0_SLEAP_UNet_topdown | 0.00678258 | 6.61E-08 | 3.25E-11 | 1 | 0.01369468 | 2.12E-06 | 0.00423038 | 0.00872865 | 1 | 1 | 1 | 1 | | | | |
| 4.0_DLC_Resnet-152 | 0.03924431 | 7.41E-07 | 4.64E-10 | 1 | 0.07368907 | 1.99E-05 | 0.02496221 | 0.04778807 | 1 | 1 | 1 | 1 | 1 | | | |
| 4.0_DLC_Resnet-50 | 1 | 1 | 0.24970976 | 0.04121778 | 1 | 1 | 1 | 1 | 0.02174299 | 0.65566065 | 1 | 0.00146362 | 0.00950779 | 1 | | |
| 4.0_SLEAP_UNet_simple | 1 | 0.000685 | 1.49E-06 | 1 | 1 | 0.00849976 | 1 | 1 | 1 | 1 | 1 | 1 | 1 | 0.90537066 | 1 | |
| 4.0_SLEAP_UNet_topdown | 1 | 1 | 0.24398669 | 0.0664869 | 1 | 1 | 1 | 1 | 0.03742037 | 0.94218088 | 1 | 0.0028804 | 0.01730841 | 1 | 1 | 1 |

**C**

**Resolution**

| Downsampled dataset | 1.0_DLC_Resnet-152 | 1.0_DLC_Resnet-50 | 1.0_SLEAP_UNet_simple | 1.0_SLEAP_UNet_topdown | 2.0_DLC_Resnet-152 | 2.0_DLC_Resnet-50 | 2.0_SLEAP_UNet_simple | 2.0_SLEAP_UNet_topdown | 3.0_DLC_Resnet-152 | 3.0_DLC_Resnet-50 | 3.0_SLEAP_UNet_simple | 3.0_SLEAP_UNet_topdown | 4.0_DLC_Resnet-152 | 4.0_DLC_Resnet-50 | 4.0_SLEAP_UNet_simple | 4.0_SLEAP_UNet_topdown |
|---|---|---|---|---|---|---|---|---|---|---|---|---|---|---|---|---|
| 1.0_DLC_Resnet-152 | 1 | | | | | | | | | | | | | | | |
| 1.0_DLC_Resnet-50 | 1 | 1 | | | | | | | | | | | | | | |
| 1.0_SLEAP_UNet_simple | 1 | 0.10399779 | 1 | | | | | | | | | | | | | |
| 1.0_SLEAP_UNet_topdown | 1 | 0.08729135 | 1 | 1 | | | | | | | | | | | | |
| 2.0_DLC_Resnet-152 | 0.12114254 | 0.00252282 | 1 | 1 | 1 | | | | | | | | | | | |
| 2.0_DLC_Resnet-50 | 1 | 0.37594502 | 1 | 1 | 1 | 1 | | | | | | | | | | |
| 2.0_SLEAP_UNet_simple | 0.05338087 | 0.00090492 | 1 | 1 | 1 | 1 | 1 | | | | | | | | | |
| 2.0_SLEAP_UNet_topdown | 1 | 0.16456702 | 1 | 1 | 1 | 1 | 1 | 1 | | | | | | | | |
| 3.0_DLC_Resnet-152 | 1.18E-05 | 3.67E-08 | 0.3630101 | 0.45108984 | 1 | 0.10003979 | 1 | 0.24632487 | 1 | | | | | | | |
| 3.0_DLC_Resnet-50 | 1.07E-07 | 1.57E-10 | 0.0201905 | 0.02664008 | 0.54254844 | 0.00419235 | 1 | 0.01262928 | 1 | 1 | | | | | | |
| 3.0_SLEAP_UNet_simple | 1 | 1 | 1 | 1 | 1 | 1 | 1 | 1 | 0.0056246 | 0.00013529 | 1 | | | | | |
| 3.0_SLEAP_UNet_topdown | 1 | 1 | 1 | 1 | 1 | 1 | 0.55775794 | 1 | 0.00040858 | 6.38E-06 | 1 | 1 | | | | |
| 4.0_DLC_Resnet-152 | 1.48E-56 | 1.48E-63 | 4.02E-41 | 1.60E-40 | 9.41E-36 | 2.04E-43 | 1.67E-34 | 1.06E-41 | 3.50E-25 | 1.53E-21 | 4.89E-48 | 1.09E-51 | 1 | | | |
| 4.0_DLC_Resnet-50 | 1.59E-21 | 5.76E-26 | 2.43E-12 | 4.58E-12 | 2.10E-09 | 1.22E-13 | 9.54E-09 | 1.05E-12 | 0.00033815 | 0.01220954 | 2.46E-16 | 1.41E-18 | 1.05E-07 | 1 | | |
| 4.0_SLEAP_UNet_simple | 3.32E-05 | 1.24E-07 | 0.66486252 | 0.81496285 | 1 | 0.19592848 | 1 | 0.46007613 | 1 | 1 | 0.01256947 | 0.00101253 | 4.27E-26 | 0.00013054 | 1 | |
| 4.0_SLEAP_UNet_topdown | 0.69503524 | 0.02350872 | 1 | 1 | 1 | 1 | 1 | 1 | 1 | 0.09296155 | 1 | 1 | 1.09E-38 | 5.22E-11 | 1 | 1 |

**D**

**Exposure**

| Downsampled dataset | 1.0_DLC_Resnet-152 | 1.0_DLC_Resnet-50 | 1.0_SLEAP_UNet_simple | 1.0_SLEAP_UNet_topdown | 2.0_DLC_Resnet-152 | 2.0_DLC_Resnet-50 | 2.0_SLEAP_UNet_simple | 2.0_SLEAP_UNet_topdown | 3.0_DLC_Resnet-152 | 3.0_DLC_Resnet-50 | 3.0_SLEAP_UNet_simple | 3.0_SLEAP_UNet_topdown | 4.0_DLC_Resnet-152 | 4.0_DLC_Resnet-50 | 4.0_SLEAP_UNet_simple | 4.0_SLEAP_UNet_topdown |
|---|---|---|---|---|---|---|---|---|---|---|---|---|---|---|---|---|
| 1.0_DLC_Resnet-152 | 1 | | | | | | | | | | | | | | | |
| 1.0_DLC_Resnet-50 | 1 | 1 | | | | | | | | | | | | | | |
| 1.0_SLEAP_UNet_simple | 1.14E-08 | 0.00026346 | 1 | | | | | | | | | | | | | |
| 1.0_SLEAP_UNet_topdown | 0.65382154 | 1 | 0.0285099 | 1 | | | | | | | | | | | | |
| 2.0_DLC_Resnet-152 | 5.80E-15 | 3.51E-09 | 1 | 2.80E-06 | 1 | | | | | | | | | | | |
| 2.0_DLC_Resnet-50 | 1.03E-11 | 1.27E-06 | 1 | 0.00038269 | 1 | 1 | | | | | | | | | | |
| 2.0_SLEAP_UNet_simple | 2.30E-11 | 2.37E-06 | 1 | 0.00063786 | 1 | 1 | 1 | | | | | | | | | |
| 2.0_SLEAP_UNet_topdown | 9.17E-40 | 4.05E-30 | 1.76E-10 | 8.76E-25 | 2.93E-05 | 1.34E-07 | 6.85E-08 | 1 | | | | | | | | |
| 3.0_DLC_Resnet-152 | 1 | 0.01779014 | 1.78E-15 | 0.00016685 | 1.87E-23 | 2.19E-19 | 6.07E-19 | 7.92E-53 | 1 | | | | | | | |
| 3.0_DLC_Resnet-50 | 1 | 1 | 1.75E-07 | 1 | 1.99E-13 | 2.39E-10 | 5.11E-10 | 2.78E-37 | 1 | 1 | | | | | | |
| 3.0_SLEAP_UNet_simple | 1 | 1 | 3.09E-06 | 1 | 8.66E-12 | 6.71E-09 | 1.36E-08 | 1.42E-34 | 0.36866078 | 1 | 1 | | | | | |
| 3.0_SLEAP_UNet_topdown | 1 | 1 | 3.05E-10 | 0.11451275 | 5.76E-17 | 1.66E-13 | 3.92E-13 | 6.16E-43 | 1 | 1 | 1 | 1 | | | | |
| 4.0_DLC_Resnet-152 | 1 | 0.15294047 | 4.95E-14 | 0.00210347 | 5.24E-22 | 6.33E-18 | 1.75E-17 | 5.69E-52 | 1 | 1 | 1 | 1 | 1 | | | |
| 4.0_DLC_Resnet-50 | 1 | 1 | 0.0037751 | 1 | 9.25E-08 | 2.58E-05 | 4.64E-05 | 1.45E-28 | 0.00033134 | 1 | 1 | 0.23015358 | 0.00426889 | 1 | | |
| 4.0_SLEAP_UNet_simple | 1 | 1 | 0.00236305 | 1 | 4.73E-08 | 1.45E-05 | 2.65E-05 | 4.23E-29 | 0.00055235 | 1 | 1 | 0.32724785 | 0.00685273 | 1 | 1 | |
| 4.0_SLEAP_UNet_topdown | 2.08E-05 | 0.07314408 | 1 | 1 | 0.06846152 | 1 | 1 | 1.97E-16 | 2.41E-11 | 0.00020457 | 0.00215614 | 9.54E-07 | 6.06E-10 | 0.63583861 | 0.45090765 | 1 |

**Colormap**

| Colormap |
|---|
| 0.05 |
| 0.04 |
| 0.03 |
| 0.02 |
| 0.01 |
| 0.008 |
| 0.006 |
| 0.004 |
| 0.002 |
| 0.001 |
| 0.0008 |
| 0.0006 |
| 0.0004 |
| 0.0002 |
| 0.0001 |
| 0.00005 |
| 0 |
